# An active learning framework improves tumor variant interpretation

**DOI:** 10.1101/2021.11.08.467747

**Authors:** Alexandra M. Blee, Bian Li, Turner Pecen, Jens Meiler, Zachary D. Nagel, John A. Capra, Walter J. Chazin

## Abstract

For precision medicine to reach its full potential for treatment of cancer and other diseases, protein variant effect prediction tools are needed that characterize variants of unknown significance (VUS) in a patient’s genome with respect to their likelihood to influence treatment response and outcomes. However, the performance of most variant prediction tools is limited by the difficulty of acquiring sufficient training and validation data. To overcome these limitations, we applied an iterative active learning approach starting from available biochemical, evolutionary, and functional annotations. The potential of active learning to improve variant interpretation was first demonstrated by applying it to synthetic and deep mutational scanning (DMS) datasets for four cancer-relevant proteins. We then probed its utility to guide interpretation and functional validation of tumor VUS in a potential biomarker for cancer therapy sensitivity, the nucleotide excision repair (NER) protein Xeroderma Pigmentosum Complementation Group A (XPA). A quantitative high-throughput cell-based NER activity assay, fluorescence-based multiplex flow-cytometric host cell reactivation (FM-HCR), was used to validate XPA VUS selected by the active learning strategy. In all cases, selecting VUS for validation by active learning yielded an improvement in performance over traditional learning. These analyses suggest that active learning is well-suited to significantly improve interpretation of VUS and cancer patient genomes.

## Introduction

Sequence-based genetic variant interpretation is a fundamental component of the study of human disease, diagnosis of genetic disorders, selection of treatments, and prediction of patient outcomes (1). In particular, precision medicine approaches to interpret variants of unknown significance (VUS) in tumors and guide clinical decision-making represent significant interests of the National Cancer Institute (NCI) (2). However, the performance of sequence-based predictive tools is limited by difficulty in acquiring sufficient benchmarking data from diverse populations and environments and a resulting lack of functional validation (3). These tools often also fail to provide specific hypotheses for mechanisms of dysfunction, which can inform predictive power and treatment selection in precision medicine.

An increasing number of rare, nonrecurrent VUS are being identified throughout tumor genomes. Interpretation of these VUS poses a significant challenge compared to recurrent hotspot variants. Rare, nonrecurrent VUS are unlikely to be the main drivers of tumor formation, but they have potential to influence progression and response to therapy. Hence, taking such VUS into account when designing a therapy can be critical to clinical outcome. Existing approaches to analyze VUS such as genome-wide association studies (GWAS) and large-scale pooled functional screens are infeasible for all genes and novel variants of interest. GWAS studies in particular have limited power for rare VUS, fail to predict the effects of single VUS of interest, cannot identify causality for single VUS, and require significant experimental follow-up (4). This represents a significant challenge for identifying reproducible, reliable biomarkers with clinical utility (5). The National Human Genome Research Institute, the American College of Medical Genetics and Genomics, and the Association for Molecular Pathology have emphasized the need for strategies that prioritize VUS for in-depth study using benchmarked, well-controlled, physiologically relevant validation assays (3,6).

The variant interpretation challenge posed by rare tumor VUS is illustrated by the reported correlation between nucleotide excision repair (NER) activity and tumor sensitivity to cisplatin treatment (7,8). NER is the primary repair mechanism for bulky DNA adducts such as those introduced by ultraviolet (UV) light and platinum (Pt)-based chemotherapeutics like cisplatin (9). Defective NER resulting from nonrecurrent VUS in Excision Repair Cross Complementation Group 2 (*ERCC2*) or from loss of *ERCC1* sensitizes tumor cells to cisplatin and leads to improved patient outcomes (10-13). In addition, recent study of The Cancer Genome Atlas (TCGA) Pan-Cancer Atlas has revealed that most genetic lesions in NER genes are nonrecurrent nonsynonymous single nucleotide variants (SNVs) with unknown impact on therapy sensitivity and cancer patient outcomes (14). Based on the studies of ERCC2 tumor VUS (11,12), a subset of the tumor VUS in other NER genes is expected to impact tumor cell response to cisplatin and other Pt-based chemotherapeutics. However, because NER genes are not known tumor drivers and there are few if any recurrent hotspot tumor mutations, NER variant interpretation is challenging.

In this report we implement an active machine learning approach to predict the NER capacity of VUS in Xeroderma Pigmentosum Complementation Group A (XPA), an essential scaffolding protein in NER (9,15-17). Germline mutations in *XPA* result in loss of NER and lead to severe phenotypes in patients with inherited Xeroderma Pigmentosum (XP) disorder including increased sensitivity to sunlight, predisposition to skin cancer, and neurological impairment (18-20). Well over 100 unique XPA VUS have been reported in tumor databases to date. These XPA tumor VUS represent an unstudied pool of variants hypothesized to measurably impact NER activity and response to Pt-based chemotherapeutics.

Machine learning paired with iterative functional validation is a promising strategy to overcome variant interpretation limitations and rapidly provide accurate annotations for VUS from tumor genomes without exhausting limited time and resources (1,21). Specifically, in an active learning strategy, VUS that are most challenging to classify by an initial machine learning model, i.e. VUS closest to the decision boundary, are functionally tested and reincorporated with new phenotypic labels in subsequent iterations of algorithm training (22,23). The approach was first benchmarked with simulations on synthetic data and available deep mutational scanning (DMS) data for four cancer-relevant proteins, using a logistic regression model trained to predict VUS effect using available biochemical, evolutionary, and functional annotations during training. We then applied this overall approach to predict the NER capacity of tumor VUS in XPA, using a limited number of labeled NER-deficient and -proficient XPA variants and unlabeled XPA VUS from tumor genomic databases. The performance of active learning was compared to traditional learning using the XPA dataset by incorporating new variant labels after measuring NER activity using a fluorescence-based multiplex flow-cytometric host cell reactivation (FM-HCR) assay. In agreement with the synthetic and DMS simulations, active learning using new NER-proficient or -deficient labels derived from FM-HCR improved algorithm performance more than traditional learning. These results establish active learning as a promising framework for overcoming limited or biased VUS training data and maximizing the utility of VUS selected for experimental evaluation.

## Materials and Methods

### Simulating active learning with synthetic and deep mutational scanning data

Synthetic data were generated from two Gaussian distributions centered at [-1, 0, 0] and [1, 0, 0] with a covariance matrix of [[1, 0, 0], [0, 1, 0], [0, 0, 1]]. Scenarios were simulated where the class distribution was balanced with a 1:1 ratio or skewed with a class ratio of 1:5. In each case, the total number of instances was 600. Deep mutational scanning (DMS) data were acquired for four proteins relevant to cancer (PTEN, TPMT, NUDT15, CYP2C9) for which variant effect on protein cellular abundance was assayed using variant abundance by massively parallel sequencing (VAMP-seq) (**Supplementary Table S1**) (24-26). Features to classify variants in the DMS proteins were compiled from the existing Database of Human Nonsynonymous SNPs and their Functional Predictions (dbNSFP) (27); 19 scores were considered encompassing physical and biochemical properties of amino acid sidechains, sequence homology, evolutionary sequence conservation, computational pathogenicity metrics based on protein stability, protein secondary structure elements, and disease-association, as well as ensemble predictors.

In each simulation experiment, training was initiated with ten labeled synthetic instances or DMS variants, either with balanced or skewed class ratios to reflect real-world scenarios. Held-out test sets were created using 10% of each dataset and maintaining the same class ratio as the overall class ratio for each to evaluate the performance of the models during each training iteration. A logistic regression model was trained on this initial dataset and the model was used to make predictions on instances in the unlabeled pool.

In the active learning approach, the five most uncertain predictions (with predicted class probabilities closest to 0.5) were selected, labeled, and added to the pool of labeled instances or variants. In the traditional learning approach, five instances or DMS variants were selected randomly, labeled, and added to the labeled pool. The logistic regression model was retrained using the updated labeled pool. This procedure was iterated 20 times to monitor the evolution of model performance as more labeled instances were added following the two different active and traditional learning strategies. Model performance was measured by the F_1_ score on the held-out test sets:

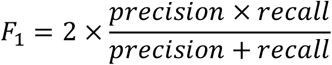

where

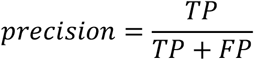

and

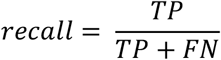

and TP: number of true positives (low-abundance variants); FP: number of false positives (wild-type like variants predicted to be low-abundance); FN: number of false negatives (low-abundance variants predicted to be wild-type like). The F_1_ score was selected because this score accounts for both precision and recall and maintains a balance between them. Because both precision and recall must be high for the final F_1_ score to be high, this metric is well-suited for variant datasets that usually exhibit an imbalance between the number of samples in each class.

### Training a logistic regression model to predict NER activity of XPA VUS

XPA variants were curated from published literature and tumor genomics databases: The NCI Genomic Data Commons Pan-Cancer Atlas, cBioPortal for Cancer Genomics, the Catalogue of Somatic Mutations in Cancer (COSMIC) v90, the Cancer Cell Line Encyclopedia (CCLE), AACR Project GENIE v7.0, and the International Cancer Genome Consortium (ICGC) data release 28. The final set of 73 tumor VUS curated from available genomics databases included only somatic single nucleotide variants (SNVs) from unique tumor samples. An additional 16 VUS were curated from the literature and were either reported without cell survival or cell-based repair activity data or had conflicting reports between studies. All 19 variants labeled as NER-proficient or NER-deficient were labeled based on reported cell survival after UV treatment or cell-based NER activity data.

Each variant was encoded with a set of 19 features that consisted of evolutionary metrics and variant scores generated by pre-existing variant pathogenicity predictors. As for the DMS simulations, these features were accessed from the Database of Human Nonsynonymous SNPs and their Functional Predictions (dbNSFP) v4.0a (27). All variants analyzed in this study and the associated references and reported data are provided in **Supplementary Tables S2** and **S3**. XPA is listed under UniProt ID: P23025; RefSeq (RRID:SCR_003496) accession number: NM_000380.3.

As several features are highly correlated (**Supplementary Figure S1**), a principal component analysis (PCA) of the feature matrix was performed (**Supplementary Figure S2**). The first three principal components were used as input features of the logistic regression model considering that the initial training set is usually very small. The model was developed using the implementation in the scikit-learn machine-learning framework (RRID:SCR_002577) (28).

The use of a semi-supervised learning algorithm was also explored to predict the NER activity of XPA VUS. A popular approach to semi-supervised learning is to create a graph that connects training instances based on their pairwise distances in the input space. Known labels are then propagated through the edges of the graph to predict the labels of unlabeled instances (29). This approach has the advantage of simultaneously using both labeled and unlabeled instances during training, compared to supervised learning algorithms. A semi-supervised label spreading model (30) was trained with the same XPA variant feature matrix used to train the logistic regression model, implemented in the scikit-learn machine-learning framework (28). The KNN kernel was used with 7 neighbors.

### Logistic regression XPA variant effect predictor with active learning and statistical analyses to compare against traditional learning

The initial logistic regression model was trained for XPA variant effect classification with the 19 variants noted above, labeled according to NER activity reported in the literature. To apply the active learning strategy to XPA, this initial model was first used to predict the class probabilities of the remaining VUS in the dataset. For the top ten VUS with the least certain predictions, i.e., probabilities closest to 0.5, (L138R, R207G, H242L, D70H, E111A, R227W, M98I, D154A, T125A, E106G, ordered from least to more certain), NER activity was measured by FM-HCR for seven VUS (L138R, H242L, D70H, E111A, D154A, T125A, E106G). In the FM-HCR analysis, VUS with NER activity significantly lower than that of wild-type XPA, with *p* < 0.05 by unpaired t tests, were labeled NER-deficient. Labeling of these assayed variants was blinded from their class probabilities predicted by the logistic regression model. To test the hypothesis that active learning improves the performance of XPA variant effect prediction more than traditional learning, a logistic regression model was retrained using a training set consisting of the initial 19 labeled variants plus the seven VUS the initial model was least certain about, labeled according to their NER activity. This was termed the “active model”.

In parallel, the NER activity was measured by FM-HCR for an additional set of 20 VUS consisting of (i) variants well separated in the PCA scatter plots and (ii) variants located in the region where the two classes are believed to overlap (**Supplementary Figure S2**). A logistic regression model was then trained using a training set consisting of the initial 19 labeled variants plus seven variants randomly selected from the pool of seven original and 20 new FM-HCR assayed variants. This was termed the “traditional model”. Next, the active and traditional model performances as measured by F_1_ scores were compared for the remaining FM-HCR assayed variants that weren’t selected for training. Due to the stochasticity in selecting variants to train the traditional model, the procedure was repeated 100 times. To enable a fair comparison, the performances of the active and traditional models were computed based on the same evaluation set in each iteration. A Mann Whitney U test was performed to compare the differences between the active and learning model performances.

### Full-length XPA model

XPA is a modular protein with two unordered regions at the N- and C-termini, which precludes an accurate representation of the 3D structure of the full-length protein in a single image. To display VUS predictions in the context of the XPA protein structure, a structural model of full-length XPA was generated based on reported XPA structures and integrative models (31-35). Starting with the coordinates of the globular XPA DNA binding domain (residues 98-239, PDBDEV00000039) (32), Rosetta FloppyTail (36) was used to model the flexible regions of XPA spanning residues 1-97 and 240-273. Default settings were used except that the perturbation cycles and models sampled parameters were increased to 1000 and 10 for each floppy tail, respectively. Graphical representations and images were generated using PyMOL Molecular Graphics System, version 2.0.7, Schrödinger, LLC (RRID:SCR_000305).

### Cell lines and cell culture

XP2OS cells (RRID:CVCL_3242) were kindly provided by Dr. Orlando Schärer (Center for Genomic Integrity, Institute for Basic Science, Ulsan National Institute of Science and Technology, Korea). Cells were maintained in DMEM (Thermo Fisher Scientific #11995073) supplemented with 10% FBS (Thermo Fisher Scientific #A3160502) and 1% Penicillin-Streptomycin (Thermo Fisher Scientific #15140122). No mycoplasma contamination was detected in this cell line throughout the experiments (SouthernBiotech #13100-01). XPA expression plasmids contain full-length human XPA (NM_000380) with the indicated mutations in the pcDNA3.1(+) backbone (GenScript custom order).

### FM-HCR assay

Reporter plasmids were prepared as a cocktail containing pMax_GFP plasmid damaged with 800 J/cm^2^ UVC radiation (herein referred to as pMax_GFP_UV) and an undamaged pMax_BFP control. An undamaged cocktail containing pMax_GFP and pMax_BFP was also utilized as a positive control. XP2OS cells (RRID:CVCL_3242) were harvested by trypsinization and pelleted via centrifugation. Cell pellets were washed with DPBS (Thermo Fisher Scientific #14190-144) and resuspended in DMEM (Thermo Fisher Scientific #11995073) supplemented with 10% FBS (Thermo Fisher Scientific #A3160502) to a final density of 2 × 10^6^ cells/mL. XP2OS cells were transfected with 200 ng of plasmid containing the XPA VUS of interest or wild-type XPA as well as the FM-HCR reporter plasmids using the Gene Pulser MXCell Plate Electroporation System (Bio-Rad Laboratories #165-2670). Plate electroporation was performed at 260 V, 950 μF.

FM-HCR analyses were performed as previously described (37,38). Briefly, fluorescence was measured via an Attune NxT Flow Cytometer (Thermo Fisher Scientific). Percent reporter expression values representing the NER capacity of cells transiently transfected with plasmids encoding each XPA variant were determined as previously described (37,38) and normalized to the NER capacity of wild-type XPA. Unpaired t-tests were performed for each wild-type and XPA variant pair (n = 3 biological replicates) using GraphPad Prism 9 (RRID:SCR_002798).

### Data Availability

The data generated in this study are available within the article and its supplementary files. All code files are available as Jupyter Notebooks in the supplement with accompanying source data.

## Results

### Active learning improves variant effect predictions for proteins with diverse functions

Active learning is a machine learning approach that incorporates iterative rounds of label determination (e.g., assigning a property from a functional assay) and training during which the algorithm chooses the data from which it learns in subsequent training rounds. Here, after functional validation of the VUS with the most uncertain initial predictions, the resulting data (e.g., variant effect on protein activity) are then used to newly label the tested variants, and the algorithm is retrained (**Figure 1**). Accurate predictions may thus be achieved using fewer rounds of training and labeling than for other strategies for validating variants (39).

**Figure 1.**
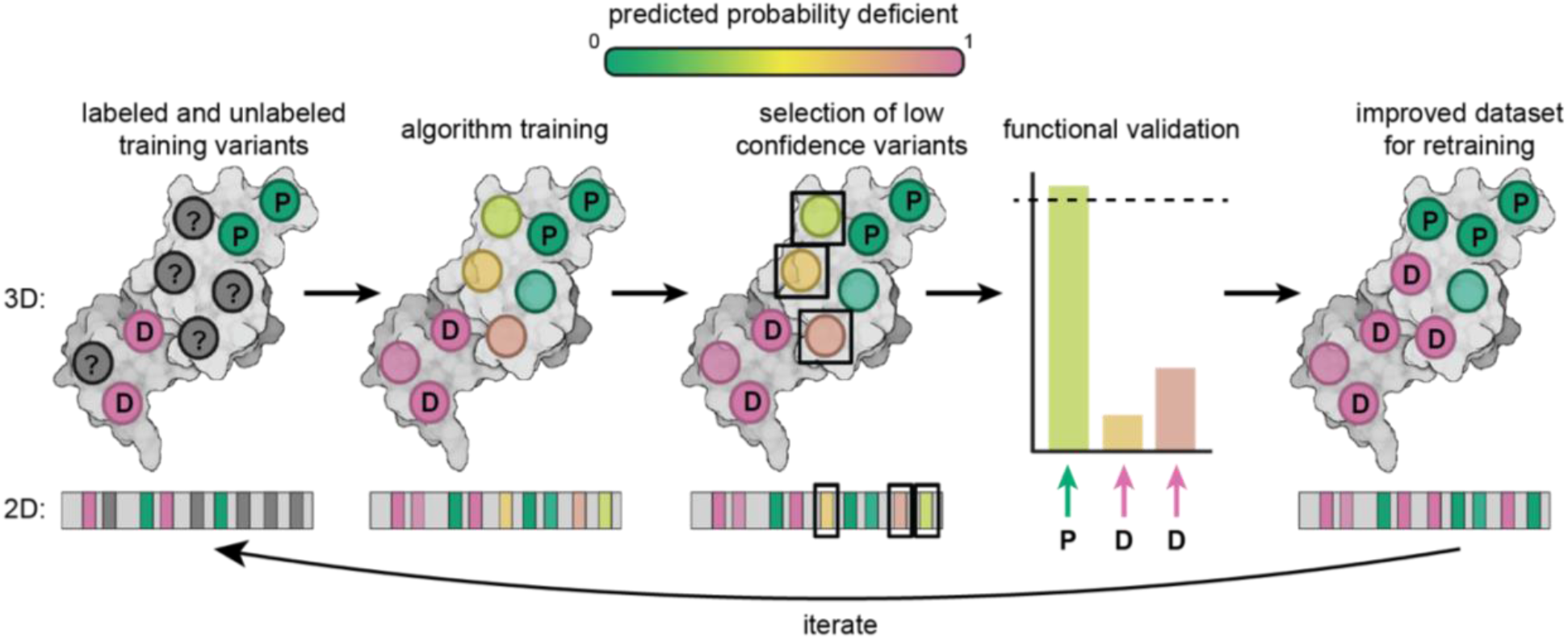
Schematic of the active learning approach to variant interpretation. First, a machine learning algorithm is trained on a set of labeled variants. Next, a subset of VUS with the lowest confidence predictions are selected and functionally validated. These newly labeled variants are then incorporated in the subsequent iteration of algorithm training. The algorithm can be retrained until predictive performance plateaus or increases only incrementally. In the diagram, NER-deficient variants are labeled with D, NER-proficient variants with P, and unlabeled VUS with a ‘?’. The color spectrum indicates the confidence of the prediction for each variant.

To test the efficacy of this proposed active learning approach before using it to guide interpretation and experimental analysis of XPA VUS, a series of simulations was performed comparing active and traditional learning on two types of data: (i) synthetic data generated from Gaussian distributions containing two binary classes of instances and (ii) real variant effect data from pre-existing DMS analyses, which quantify the effects of every possible amino acid substitution within a protein in cells and provide deleterious or neutral molecular phenotype labels for each variant. For these simulations, synthetic instances or DMS variants were present in two classes, and the identity of each synthetic instance or the phenotype associated with each DMS variant was either included as a label or excluded, resulting in unlabeled datapoints. Within the DMS analyses, we focused on four proteins with known roles in tumor suppression, progression, or therapeutic response: phosphatase and tensin homolog (PTEN) (24), thiopurine S-methyltransferase (TPMT) (24), Nudix hydrolase 15 (NUDT15) (26), and cytochrome P450 family 2 subfamily C member 9 (CYP2C9) (25). In addition to the phenotypic labels, we compiled 19 features for each DMS variant from the Database of Human Nonsynonymous SNPs and their Functional Predictions (dbNSFP) to be used as input features for training and classification (27), These features encompassed physical and biochemical properties of amino acid sidechains, sequence homology, evolutionary sequence conservation, computational pathogenicity metrics based on protein stability, protein secondary structure elements, and disease-association.

For each type of data, an uncertainty sampling query strategy (active learning) was compared to a random sampling strategy (traditional learning) (**Figure 2A**). A logistic regression model was trained for these analyses (23); we note that other algorithms could be used within the active learning framework. In a real-word scenario, the set of labeled data available for training the initial iteration of the algorithm will often come from variants previously tested and reported in the literature. Thus, the distribution of initial training data between the two possible binary classifications for each variant may not reflect the overall ratio for all possible variants in the protein. This was true for the DMS data, where each protein of interest exhibited varying ratios between the number of variants with wild-type or protein-deficient phenotypes (**Supplementary Table S1**). To reflect this reality in our simulations, differing class ratios of labeled variants were tested in the initial labeled training sets and changes in algorithm performance were measured over 20 iterations of active and traditional learning. During active learning, synthetic instances or DMS variants with the most uncertain predictions were identified and labeled based on the binary class to which they belonged.

**Figure 2.**
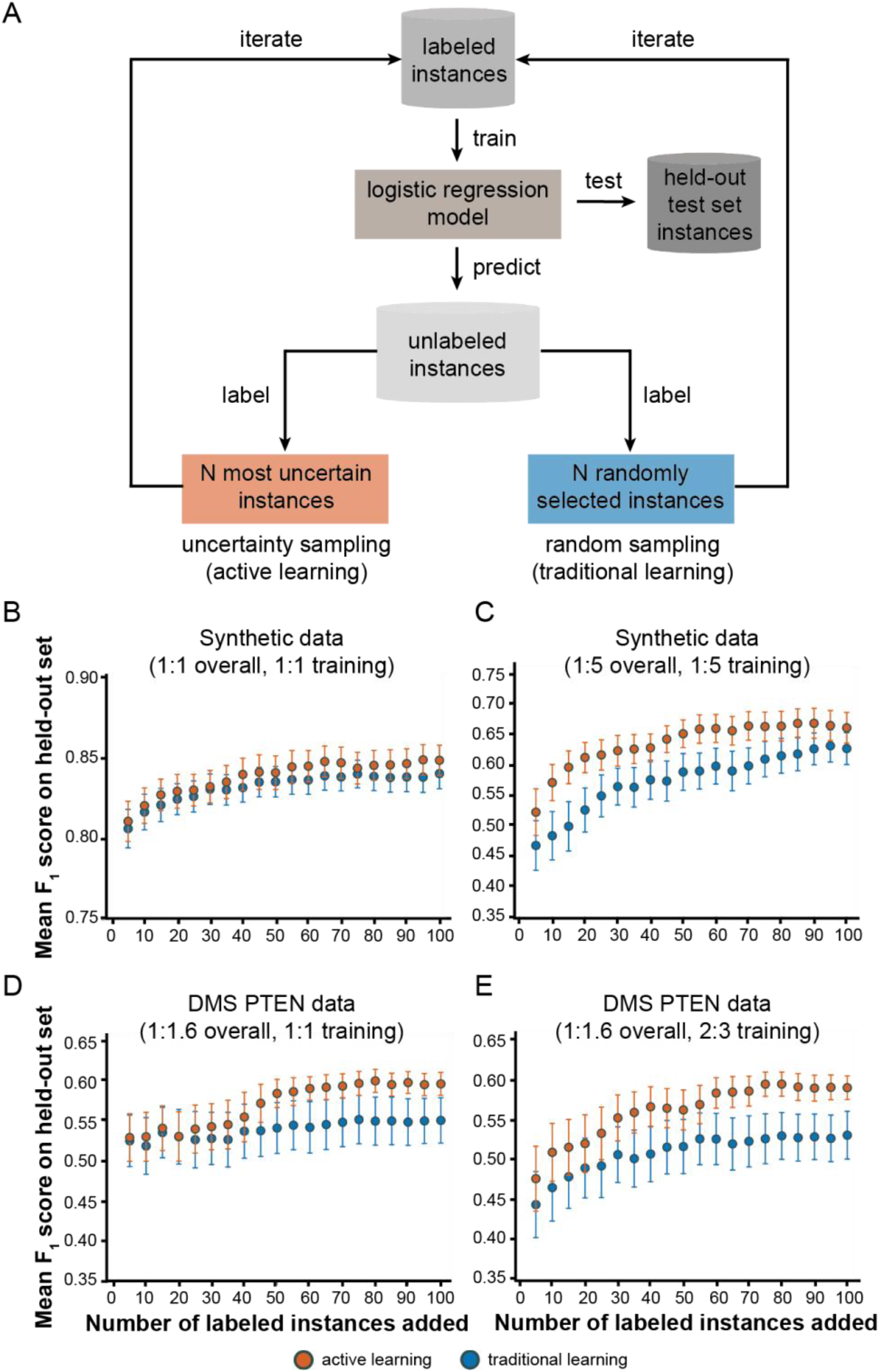
Active learning results in more accurate models compared to traditional learning on synthetic and deep mutational scanning data. *A*, Schematic representation of the simulation protocol to compare active learning with traditional learning. The mean F_1_ score was used to compare active and traditional learning for: synthetic datasets with balanced class ratios (1:1) in both the overall data and the initial labeled training set in *B*, or skewed class ratio (1:5) in both the overall data and initial labeled training set in *C*; and a DMS PTEN dataset with a balanced class ratio (1:1) in the initial labeled training set in *D*, or a skewed class ratio (2:3) in the initial labeled training set in *E*. Error bars indicate 95% confidence intervals around the mean F_1_ score. All initial labeled pools had ten instances or variants to start except for the skewed synthetic dataset in *B*, which had 12 instances to maintain the 1:5 ratio with sufficient starting numbers of instances in both classes. See **Supplementary Table S1** for additional details regarding the composition of the PTEN dataset.

Active learning achieved stronger performance than traditional learning in nearly all scenarios (**Figure 2** and **Supplementary Figure S3**). For example, in one DMS simulation, active learning outperformed traditional learning by a mean F_1_ score of 0.052 across the 20 iterations (*p* = 3.44 × 10^−14^, two-sided paired t-test) (**Figure 2E**). Similar improvement of active learning over traditional learning was achieved in all other simulations except in two exceptional scenarios. In the first, the class ratios of the initial pool of synthetic instances (5:1 or 7:3) were heavily skewed opposite to the overall class ratio of the dataset as a whole (1:5 or 1:1.9) (**Supplementary Figures S3D** and **S3H**). In the second, for CYP2C9 (**Supplementary Figures S3L-N**), active learning provided notable benefits in the early training iterations with the most limited proportions of labeled data, although this benefit decreased in later iterations as larger proportions of training data were labeled. Nevertheless, using active learning to train a variant effect predictor enabled flexible integration of pre-existing phenotypic data and reduced the time and resources needed to improve predictions. Given these primarily positive results, we next applied a similar active learning approach to XPA tumor VUS.

### Prediction of XPA VUS effects on NER

As an essential NER scaffolding protein, XPA performs two key functions during repair: (i) DNA binding at the junction between single strand and double strand DNA that is formed upon opening of the ‘repair bubble’ (15-17), and (ii) interaction with multiple proteins that constitute the NER machinery (9,32,40-43) (**Figure 3A**). Previous functional study of specific XPA variants, such as those variants known to cause the germline inherited disorder XP, were used to classify and assign labels to an initial training dataset with 19 labeled variants (8 NER-proficient and 11 NER-deficient). An additional 89 unlabeled VUS were curated primarily from publicly available tumor genomic databases to comprise the rest of the dataset (**Figure 3B**; **Supplementary Tables 2** and **3**).

**Figure 3.**
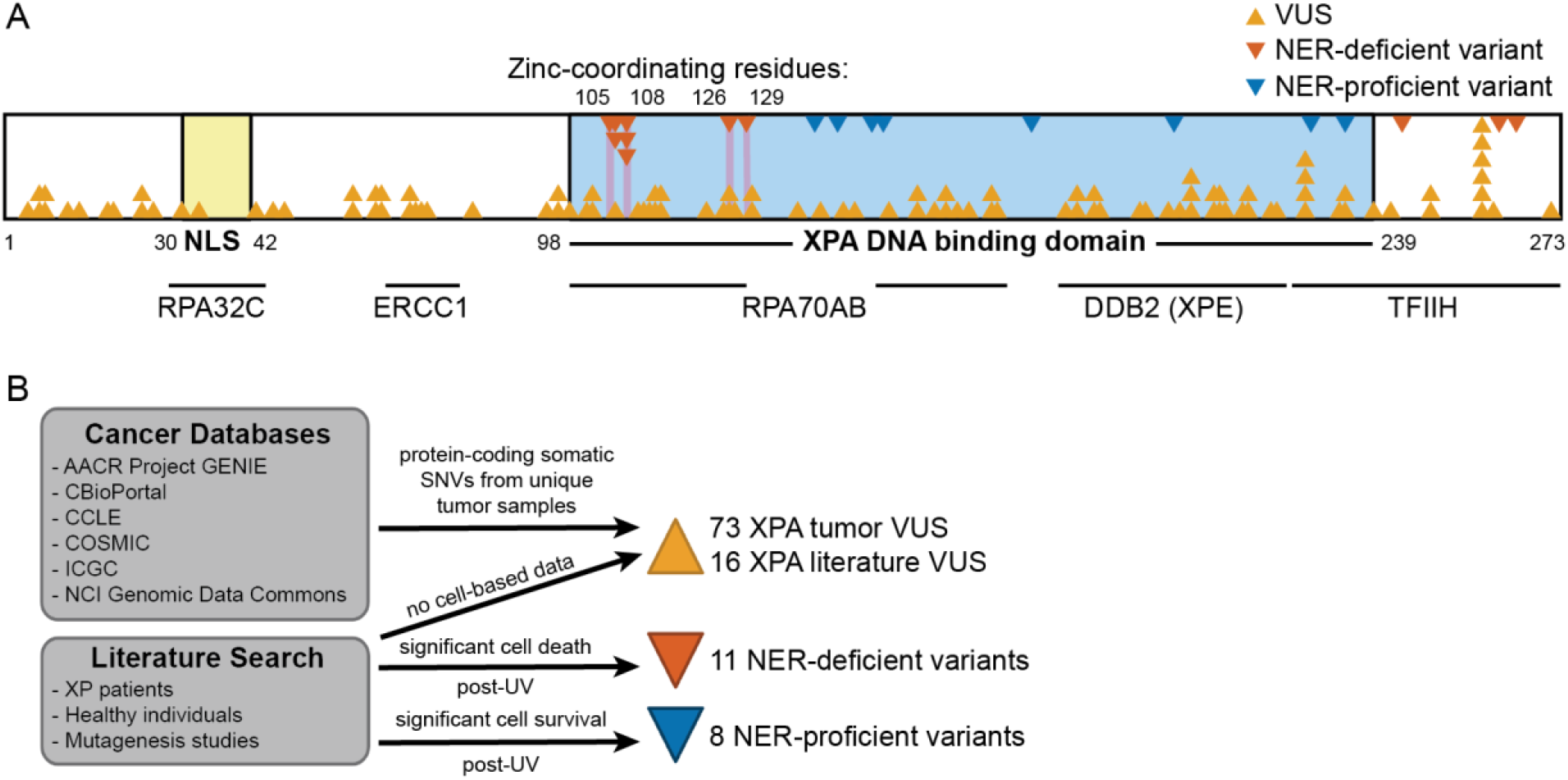
XPA contains many VUS and few functionally characterized variants. *A*, Schematic representation of the XPA protein with variants and partner protein interaction regions (horizontal lines) mapped across the sequence. The locations of NER-deficient or - proficient variants as well as VUS are indicated with triangles. *B*, Diagram outlining the sources of variants and labels used for training the initial variant effect prediction algorithm.

Following the approach used for the DMS analysis, 19 features for each XPA variant were compiled from dbNSFP including: amino acid properties, sequence homology, evolutionary sequence conservation, computational variant pathogenicity, and ensemble scores. The features exhibited substantial variability across variants (**Figure 4A**; **Supplementary Figure S1**) and inspection of the ability of these scores to distinguish known NER-deficient and -proficient XPA variants revealed clear room for improvement (**Supplementary Table 4**). These data further emphasize the need for an approach that incorporates functional data specific to the protein and phenotype of interest.

**Figure 4.**
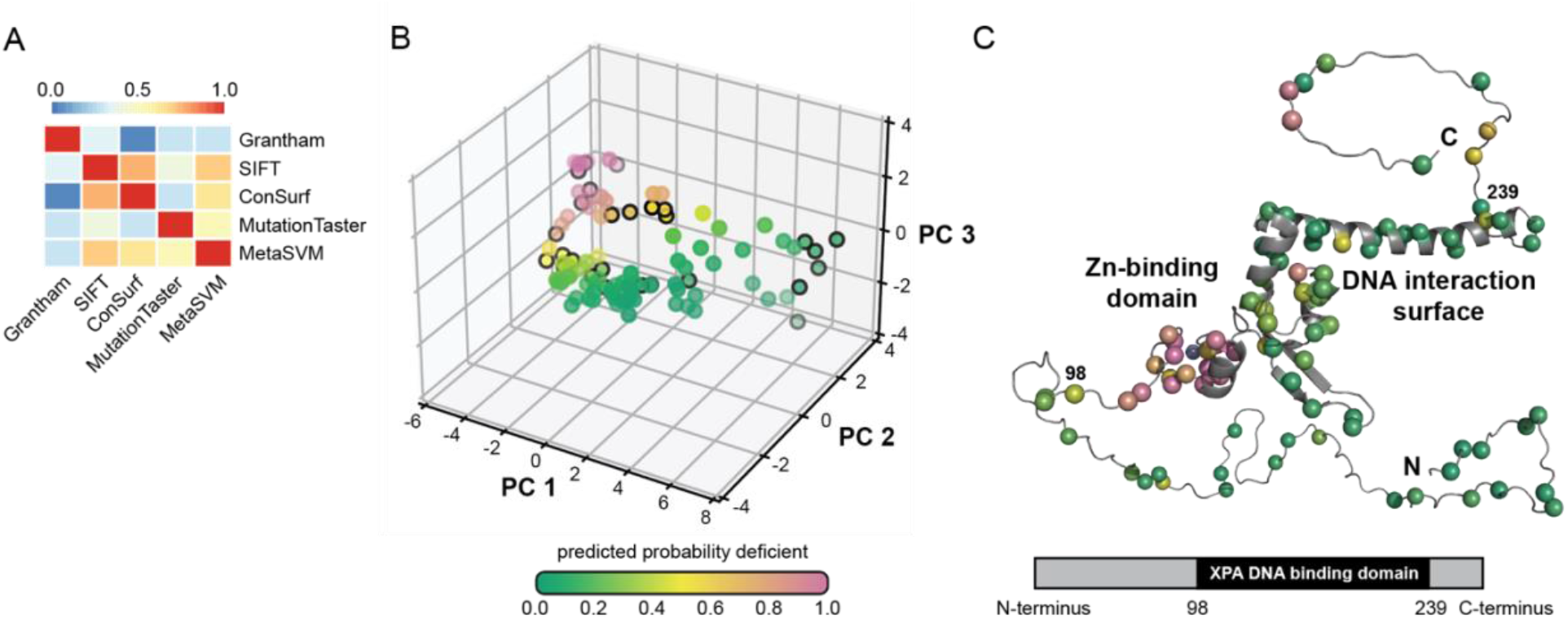
Logistic regression model to predict NER-deficient variants. *A*, Heatmap of pairwise Spearman’s rank correlations of five representative features for each XPA variant. Features shown include one predictor from each of the following classes: amino acid properties (Grantham), sequence homology (SIFT), evolutionary sequence conservation (ConSurf), pathogenicity (MutationTaster), and ensemble scores (MetaSVM). *B*, Effects of XPA VUS on NER activity predicted by the logistic regression model. Input features are the first three principal components from a principal component analysis (PCA) of the original set of 19 features from dbNSFP. VUS selected for functional validation outlined in black: D5Y, G6R, A18S, R30W, A60T, D70H, G72E, G73E, P94L, E106G, K110E, E111A, F112C, M113I, D114Y, T125A, C126W, C126Y, R130I, L138R, Y148D, D154A, F164C, V234M, H242L, R258C, and K272N. *C*, Model of full-length XPA with variants of interest depicted as spheres and colored according to the scheme in *B* (top). The precise fold and orientation of the flexible N- and C-termini regions are not known and are shown only for representative purposes. The bottom panel shows a schematic diagram of XPA and the location of the XPA DNA binding domain.

Given the limited amount of training data for XPA, the dimensionality of the initial feature set was reduced using principal component analysis (PCA) before training a logistic regression algorithm (**Supplementary Figure S2**). Mapping the initial predictions as the probability of being classified NER-deficient onto the PCA of the variant features revealed clusters of high-confidence predicted NER-proficient and -deficient variants, with a population of lower confidence predictions at the boundaries between clusters (**Figure 4B**). We also observed similar patterns when making predictions using a semi-supervised label spreading algorithm (30,44,45) to analyze the XPA dataset **(Supplementary Figure S4; Supplementary Tables S5, S6)**.

The NER-deficient class probability for each variant was mapped onto a structural model of XPA, further supporting the algorithm predictions. For example, coordination of a zinc atom by cysteine residues 105, 108, 126, and 129 is required for the structural and functional integrity of XPA (46). Hence, tumor VUS such as C126W and VUS in adjacent residues were predicted to be NER-deficient (**Figure 4C**). In contrast, mutagenesis studies have demonstrated that single mutation of residues along the large DNA binding surface of the XPA DBD are sometimes insufficient to abrogate DNA binding and NER activity (47,48), and fewer VUS on this surface were predicted to be NER-deficient (**Figure 4C**). Similarly, H244R, C261S, and C264S in the flexible C-terminus have been shown to be NER-deficient, and the nearby tumor VUS H242L was predicted to also be NER-deficient (**Figure 4C**). These results demonstrate the potential of variant effect prediction for XPA VUS.

### Active learning using functional validation improves variant effect predictions for XPA

To determine the effect of incorporating functional validation into our approach, 27 VUS were selected for functional validation by FM-HCR, a high-throughput host cell reactivation assay to quantify NER capacity (37) (**Figure 5A**). These VUS spanned the spectrum of prediction confidence, enabling evaluation of algorithm performance and comparison of active learning with traditional learning. This set included seven of the ten VUS with least certain class probabilities from the initial logistic regression model and an additional 20 VUS for evaluation of model performance.

**Figure 5.**
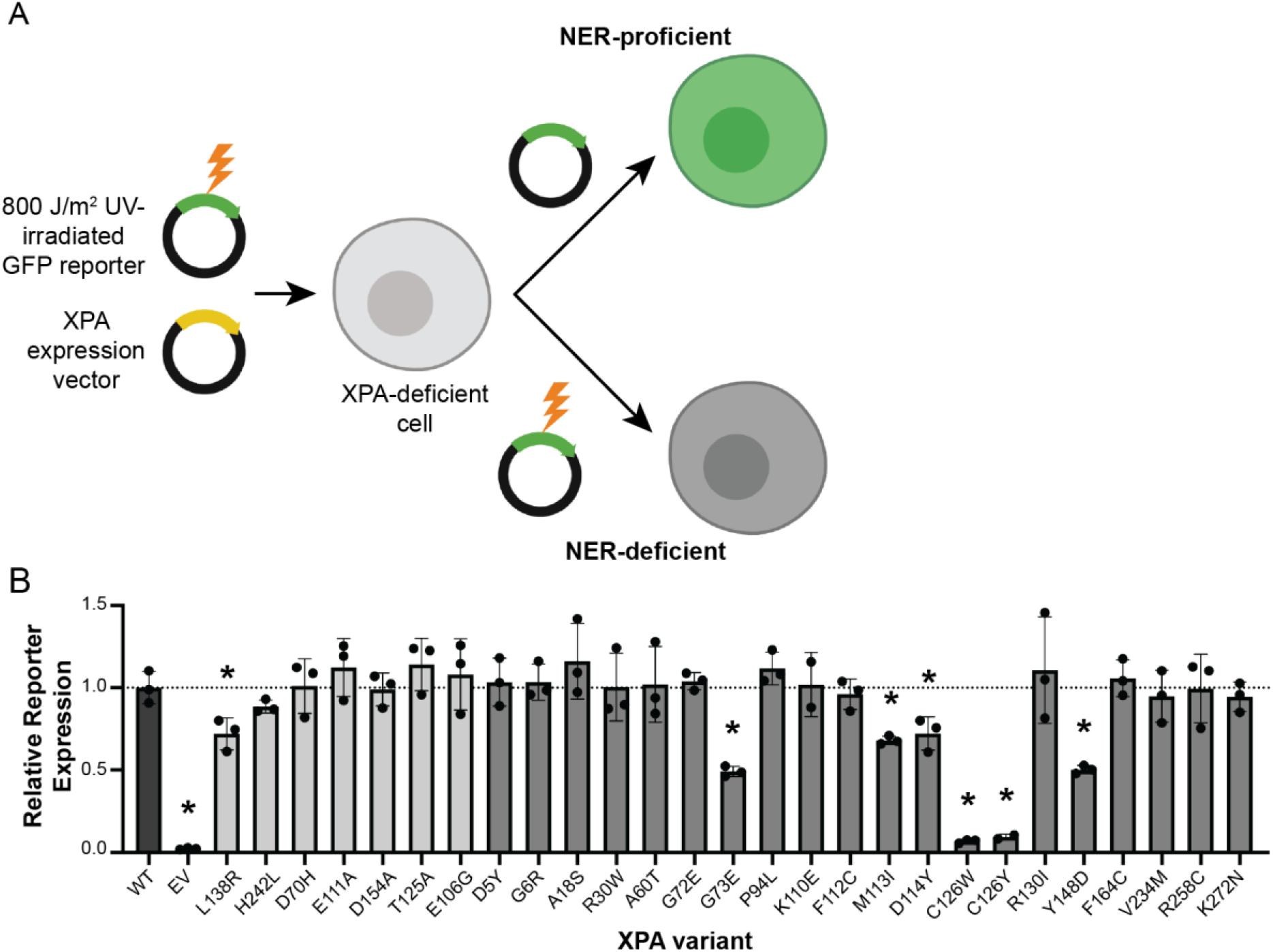
FM-HCR to test NER capacity of selected XPA VUS. *A*, Diagram of FM-HCR assay in XPA-deficient XP2OS cells. Cells transfected with UV-damaged fluorescent reporters as well as either WT XPA or XPA VUS are analyzed by flow cytometry to detect fluorescent reporter expression. Successful NER results in fluorescent reporter repair and expression (top), which is not observed in control cells lacking XPA (bottom). *B*, Bar graph showing relative reporter expression in cells expressing empty vector (EV), WT XPA, or the 27 VUS selected for validation. Seven of the top ten VUS with the least certain class probabilities (light grey) were tested, as well as 20 other VUS for further evaluation (dark grey). Damaged reporter expression was normalized to an undamaged control reporter to account for transfection efficiency. The percent reporter expression for each variant was normalized to that determined for WT to generate the final relative reporter expression (n = 3 biological replicates). Error bars indicate standard deviation from the mean. Seven of the VUS analyzed maintained significantly decreased repair capacity when compared to WT. * signifies *p* < 0.05, unpaired t test.

The XPA VUS selected for FM-HCR were transiently overexpressed in XPA-deficient XP2OS cells (49), together with a UV-damaged green fluorescent protein (GFP)-expressing reporter. Successful NER of the UV-damaged reporter in NER-proficient cells can be detected and quantified by flow cytometry (**Figure 5A**). As anticipated, XPA-deficient XP2OS cells had very little GFP reporter expression relative to XP2OS cells rescued with wild-type (WT) XPA (**Figure 5B**). Several variants rescued NER activity to a similar degree, but not significantly beyond that of WT XPA, providing assurance that cells transiently complemented with different expression constructs can achieve similar levels of NER capacity as WT (**Figure 5B**). The FM-HCR results also revealed a gradient of NER deficiency resulting from a subset of variants. As predicted, profound NER defects were observed upon substitution of residues that coordinate the zinc ion, such as C126 (**Figure 5B**). Notably, many variants predicted to be deleterious by pre-existing predictors were not associated with significant NER-deficiency and vice versa (**Supplementary Table S7**). Comparison of our initial algorithm predictions with these functional data provided the basis for an iterative active learning approach (**Supplementary Table S8**).

To further evaluate the active learning approach, the logistic regression model was retrained using 26 labeled training variants. The original 19 training set variants were used with labels assigned based on previous characterization in the literature. In addition, seven VUS from the group least confidently predicted by the initial model were added using the newly assigned NER-proficient or -deficient labels from the FM-HCR analysis. The active learning model was compared to F_1_ scores from 100 traditional learning models trained using the original 19 labeled variants plus seven variants randomly selected from the remaining 20 variants assayed by FM-HCR. To enable a fair comparison, the active learning model was evaluated on the same held-out variants as each of the 100 traditional learning models, and thus, we also obtained 100 F_1_ scores for the active learning approach. Consistent with our hypothesis, the active learning model performed significantly better than the traditional learning model (mean F_1_ score 0.752 vs. 0.650 for 100 trials, *p* = 3.8 × 10^−10^, Mann Whitney U test) (**Figure 6**). This improvement in performance illustrates that active learning is practical and beneficial in real-life situations where the amount of initial training data is small and obtaining additional labels is costly and laborious.

**Figure 6.**
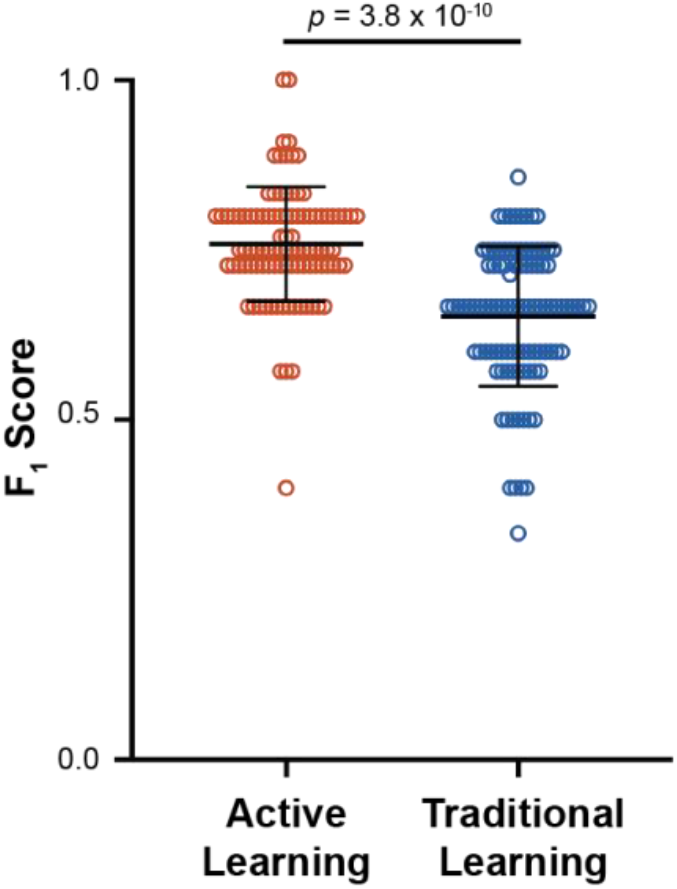
Active learning improves predictions of XPA variant NER capacity. Plot of F_1_ scores comparing the performance of active versus traditional learning with the XPA dataset. Both the active and traditional learning strategies were repeated 100 times. Error bars indicate standard deviation. *p* = 3.8 × 10^−10^, Mann Whitney U test.

## Discussion

Our analyses of synthetic, DMS, and real-world XPA variant datasets demonstrate that active learning and targeted functional validation focused on variants that are refractory to algorithmic classification can address current variant interpretation challenges. Functional validation is increasingly recognized as a centerpiece of variant interpretation (3,6,50), and active learning provides an efficient framework to guide the selection and incorporation of validation data for maximal impact. Screening out variants unlikely to be informative and prioritizing others for follow-up avoids wasted experimental effort and has the potential to more rapidly identify variants with functional effects. Our analyses provide the basis for future work to predict, screen, and conduct in-depth studies of XPA VUS that reduce NER activity and sensitize cells to cisplatin.

The analyses of synthetic and DMS data identified a few discrete examples where active learning failed to significantly improve performance compared to traditional learning. Notably, this occurred in scenarios with class ratios for the overall dataset that were heavily skewed opposite to the subset of labeled training instances (**Supplementary Figures S3D** and **S3H**). This finding reveals a limitation in how sparse or biased the initial training dataset can be while still generating accurate predictions. It also suggests that active learning cannot fully overcome severe under-representation of variant classes in the training set that are more prevalent in the overall data. However, given that the sources of labels used for training are known, it should be possible to foresee when there is likely to be a substantial ascertainment bias that could decrease the utility of active learning. The results for the CYP2C9 DMS data also hint that the success of active learning may be context dependent. While active learning showed improvement over traditional learning for CYP2C9 during the early iterations with the most limited proportion of labeled training data, which likely reflects most real-world scenarios, improvement was small in later rounds (**Supplementary Figures S3L-N**). More thorough exploration of DMS and other data will be necessary to clearly define the scenarios in which active learning is most beneficial.

We have demonstrated that active learning can be successfully applied using inputs derived from either functional data or computational predictions of functional significance to improve variant effect predictions. This is a central strength, particularly because active learning can also be easily extended to include additional phenotypic data of interest such as protein structural data and other functional assays, which would both be expected to improve predictive performance. Using phenotypic data such as drug sensitivity to validate variant labels during training represents one future area of exploration that may allow for the generalization of this approach to other proteins or protein complexes.

Improved performance of XPA variant interpretation is anticipated with higher quality and consistency of labels in the training set. The initial XPA variant training labels used here were derived from published results of different cell-based assays from various research groups and the specific variants were selected subjectively. Starting with standardized, quantifiable FM-HCR analyses to derive accurate labels for the entire initial training set is expected to greatly improve predictive performance. Future studies will include updating the active learning model by retraining with XPA variants labeled solely by high quality FM-HCR analysis and conducting several additional iterations of active learning. Incorporating deeper insights into the structure and mechanisms of the NER machinery into training is also anticipated increase the performance of VUS interpretation. This information will also enable the development of hypotheses about mechanisms of NER dysfunction, which in turn can be tested and refined using cell-based, biochemical, biophysical, and structural analysis.

Our analyses underscore that single XPA tumor VUS have the potential to abrogate NER activity in cells, irrespective of other genetic events. However, there are many VUS in NER proteins within the same tumor samples that could influence NER activity; tumor cells are complex and variant interpretation should consider all potentially relevant variants in an individual (14). Nonetheless, even with these limitations, the active learning strategy paired with FM-HCR validation shows significant promise for XPA variant interpretation. One goal on the horizon is to better understand and predict tumor cell drug sensitivity using higher performing models to identify XPA variants as biomarkers for cisplatin response. This would involve directly testing repair of cisplatin-induced lesions in cells expressing tumor VUS. Ultimately, this machine learning approach and future improved versions are anticipated to enable prediction of the cisplatin response in cells expressing a broad range of NER VUS.

Active learning can overcome small training datasets, enable the selection of a feasible number of VUS for validation, and maximize the performance gains provided by cell-based functional validation. By providing actionable insights into VUS, this approach contributes to the successful implementation of cancer precision medicine.

## Supporting information

Zipped file containing three Jupyter Notebooks with the code used in the study and five source data files required to run the Jupyter Notebooks

## Acknowledgements

We acknowledge Dr. Orlando Schärer for his generous gift of XP2OS cells, and Dr. Jonathan Sheehan and Dr. Chris Moth in the Vanderbilt Program in Personalized Structural Biology for valuable training and variant interpretation insights. Some diagrams were created with a licensed version of BioRender.com.

## Author Contributions

Conceptualization: WJC, JAC, AMB, BL

Methodology: JAC, BL, AMB, JM, ZDN, TP

Software: BL

Validation: BL, AMB, ZDN, TP

Formal Analysis: BL, AMB, JAC, ZDN, TP

Investigation: BL, AMB, TP

Data Curation: AMB, BL

Writing – Original Draft: AMB, BL, JAC, WJC

Writing – Review and Editing: JAC, WJC, AMB, BL, JM, ZDN, TP

Visualization: AMB, BL, TP

Supervision: WJC, JAC, ZDN

Funding Acquisition: WJC, JAC, ZDN, AMB

