## Supplementary material for "An active learning framework improves tumor variant interpretation": Zipped file containing three Jupyter Notebooks with the code used in the study and five source data files required to run the Jupyter Notebooks: active_learning_dms.ipynb

```
# %% [markdown]
### Exploring Active Learning Using DMS Data

# %% [code]
%reset

import os
import numpy as np
import pandas as pd
import matplotlib.pyplot as plt

# data directory
data_dir = '/path/to/where/the/features/are/located/'

# load data file
dms_file = os.path.join(data_dir, 'cyp2c9_pca_features.csv')
dms_df = pd.read_csv(dms_file)
dms_df = dms_df.drop_duplicates(['variant'])

# toggle the following cutoff based on the dataset used 
labels = []
cutoff = 0.77 # CYP2C9
# cutoff = 0.72 # NUDT15
# cutoff = 0.71 # PTEN, below this cutoff is bad
# cutoff = 0.72 # TPMT, below this cutoff is bad
for x in dms_df.dms_score:
    if x < cutoff:
        labels.append(1)
    else:
        labels.append(0)
dms_df['label'] = labels

# choose training examples randomly
dms_a_X = np.array(dms_df[dms_df.label == 1])[:, 1:4]
dms_b_X = np.array(dms_df[dms_df.label == 0])[:, 1:4]

# %% [markdown]
#### Assigned labels and extract features

# %% [code]
# import required packages
import numpy as np
import pandas as pd
from sklearn.metrics import accuracy_score, f1_score
from sklearn.linear_model import LogisticRegression
import plotly.express as px
np.random.seed(0)

# comparing the performance of active learning and traditional learning
j = 0
all_accs_active = []
all_accs_random = []

# 10% of the dataset will be set aside for testing
a_test = int(round(len(dms_a_X) * 0.1))
b_test = int(round(len(dms_b_X) * 0.1))
while j < 100:
    # set aside 10% of the data for testing
    pmt_idx_a = np.random.permutation(len(dms_a_X))
    pmt_idx_b = np.random.permutation(len(dms_b_X))
    test_a_X = dms_a_X[pmt_idx_a[:a_test]]
    train_a_X = dms_a_X[pmt_idx_a[a_test:]]
    test_b_X = dms_b_X[pmt_idx_b[:b_test]]
    train_b_X = dms_b_X[pmt_idx_b[b_test:]]

    test_X = np.concatenate((test_a_X, test_b_X), axis=0)
    test_y = np.concatenate((np.repeat(1, a_test), np.repeat(0, b_test)), axis=0)

    # initial training data
    # these two numbers will need to based updated when a new dataset is used
    # n_init_pos + n_init_neg should be equal to 10, and n_init_pos:n_init_neg should be 
    # equal to the class ratio of the overall dataset
    n_init_pos = 6
    n_init_neg = 4
    n_pos  = len(train_a_X)
    n_neg = len(train_b_X)
    X_init_a = train_a_X[:n_init_pos]
    X_init_b = train_b_X[:n_init_neg]
    X_init = np.concatenate((X_init_a, X_init_b), axis=0)
    y_init = np.concatenate((np.repeat(1, n_init_pos), np.repeat(0, n_init_neg)), axis=0)

    X_rest = np.concatenate((train_a_X[n_init_pos:], train_b_X[n_init_neg:]), axis=0)
    y_rest = np.concatenate((np.repeat(1, n_pos - n_init_pos), np.repeat(0, n_neg - n_init_neg)), axis=0)

    # train an initial model
    clf_model = LogisticRegression()
    clf_model.fit(X_init, y_init)

    # make predictions on the rest of the dataset
    pred_probs = clf_model.predict_proba(X_rest)[:, 0]

    # active learning
    def get_least_confident_idx(pred_prob, n=5):
        idx_sorted = np.argsort(np.abs(pred_prob - 0.5))
        return idx_sorted[:n], idx_sorted[n:] 

    accs_active = []
    i = 0
    X_combined_active = X_init
    y_combined_active = y_init
    X_rest_active = X_rest
    y_rest_active = y_rest
    while i < 20:
        # shuffled_indices = np.random.permutation(unsure_idx)
        new_idx, rest_idx = get_least_confident_idx(pred_probs, n=5)
        X_new_active = X_rest_active[new_idx]
        y_new_active = y_rest_active[new_idx]

        # label spreading model
        X_combined_active = np.concatenate((X_combined_active, X_new_active), axis=0)
        y_combined_active = np.concatenate((y_combined_active, y_new_active), axis=0)
        clf_model_active = LogisticRegression()
        clf_model_active.fit(X_combined_active, y_combined_active)    

        # predict labels on test set
        pred_labels = clf_model_active.predict(test_X)
        accs_active.append(f1_score(test_y, pred_labels))

        # update the remaining data
        X_rest_active = X_rest_active[rest_idx]
        y_rest_active = y_rest_active[rest_idx]

        # predict probabilities for the remaining data
        pred_probs = clf_model_active.predict_proba(X_rest_active)[:, 0]

        # increment i
        i = i + 1
    all_accs_active.append(accs_active)
    
    # traditional learning
    accs_random = []
    k = 0
    n = 5
    X_combined_random = X_init
    y_combined_random = y_init
    X_rest_random = X_rest
    y_rest_random = y_rest
    while k < 20:
        # features
        shuffled_indices = np.random.permutation(len(X_rest_random))
        X_new_random = X_rest_random[shuffled_indices[:n]]
        y_new_random = y_rest_random[shuffled_indices[:n]]
        X_combined_random = np.concatenate((X_combined_random, X_new_random), axis=0)
        y_combined_random = np.concatenate((y_combined_random, y_new_random), axis=0)

        # label spreading model
        clf_model_random = LogisticRegression()
        clf_model_random.fit(X_combined_random, y_combined_random)    

        pred_labels = clf_model_random.predict(test_X)
        accs_random.append(f1_score(test_y, pred_labels))

        # update the remaining data
        X_rest_random = X_rest_random[shuffled_indices[n:]]
        y_rest_random = y_rest_random[shuffled_indices[n:]]

        # increment k
        k += 1
    all_accs_random.append(accs_random)
    
    j += 1

# %% [code]
# import required packages
import numpy as np
import pandas as pd
import scipy.stats as st
import plotly.express as px

def compute_ci(data, level=0.05):
    ci = []
    for i in range(data.shape[1]):
        ci.append(
            st.t.interval(1 - level, data.shape[0] - 1, loc=np.mean(data[:, i]), scale=st.sem(data[:, i]))
        )
    return np.array(ci)

random_arr = np.array([
    np.arange(5, 105, 5) , 
    np.mean(all_accs_random, axis=0), 
    np.std(all_accs_random, axis=0),
    compute_ci(np.array(all_accs_random))[:, 0],
    compute_ci(np.array(all_accs_random))[:, 1]
])
active_arr = np.array([
    np.arange(5, 105, 5) , 
    np.mean(all_accs_active, axis=0), 
    np.std(all_accs_active, axis=0),
    compute_ci(np.array(all_accs_active))[:, 0],
    compute_ci(np.array(all_accs_active))[:, 1]    
])
arr_combined = np.concatenate((random_arr.T, active_arr.T), axis=0)

df = pd.DataFrame(arr_combined, columns=['n_train', 'mean_accuracy', 'std', 'ci_low', 'ci_high'])
df['e_minus'] = df['mean_accuracy'] - df['ci_low']
df['e'] = df['ci_high'] - df['mean_accuracy']

df.loc[:, 'strategy'] = 'random'
df.loc[20:, 'strategy'] = 'active'

fig = px.scatter(
    df, x='n_train', y='mean_accuracy',
    error_y_minus='e_minus',
    error_y='e',
    color='strategy',
    color_discrete_sequence=['#0172B2', '#D55E00'],
    labels={
        'n_train': 'Number of labeled variants added',
        'mean_accuracy': 'Mean F1 score on held-out set'
    },
    range_y=[0.60, 0.80]
)
fig.update_traces(
    marker=dict(
        size=12,
            line=dict(
            width=2,
            color='DarkSlateGrey'
        )
    ),
    selector=dict(mode='markers')
)
fig.update_layout(
    font_family='Arial',
    font_size=18,
    showlegend=False,
    xaxis=dict(
        tickmode='linear',
        tick0=10,
        dtick=10,
        ticks='outside'
    ),
    yaxis=dict(
        tickmode='linear',
        tick0=0.55,
        dtick=0.05,
        ticks='outside',
        tickformat='.2f'
    )
)
fig.update_layout({
    'plot_bgcolor': 'rgba(0, 0, 0, 0)',
})

fig.show()
```
