## Supplementary material for "An active learning framework improves tumor variant interpretation": Zipped file containing three Jupyter Notebooks with the code used in the study and five source data files required to run the Jupyter Notebooks: active_learning_synthetic_data.ipynb

```
# %% [code]
%reset
import numpy as np
from sklearn.metrics import accuracy_score, f1_score
from sklearn.linear_model import LogisticRegression

# %% [code]
# set random seed so that the result is reproducible
np.random.seed(0)

def get_least_confident_idx(pred_prob, n=5):
    """
    Returns the indices of the top 5 least confidence instances.
    """
    idx_sorted = np.argsort(np.abs(pred_prob - 0.5))
    return idx_sorted[:n], idx_sorted[n:] 

# synthetic data, two classes a and b
cov = 1
syn_a_X = np.random.multivariate_normal([-1, 0, 0], [[cov, 0, 0], [0, cov, 0], [0, 0, cov]], size=300)
syn_b_X = np.random.multivariate_normal([1, 0, 0], [[cov, 0, 0], [0, cov, 0], [0, 0, cov]], size=300)

# comparing the performance of active learning and traditional learning
i = 0 
all_accs_active = []
all_accs_random = []
a_test = 30
b_test = 30
while i < 100:
    # set aside 10% of the data for testing
    pmt_idx_a = np.random.permutation(len(syn_a_X))
    pmt_idx_b = np.random.permutation(len(syn_b_X))
    test_a_X = syn_a_X[pmt_idx_a[:a_test]]
    train_a_X = syn_a_X[pmt_idx_a[a_test:]]
    test_b_X = syn_b_X[pmt_idx_b[:b_test]]
    train_b_X = syn_b_X[pmt_idx_b[b_test:]]

    # class a is labeled 1, class b is labeled 0
    test_X = np.concatenate((test_a_X, test_b_X), axis=0)
    test_y = np.concatenate((np.repeat(1, a_test), np.repeat(0, b_test)), axis=0)

    # initial training data
    n_init_a = 6
    n_init_b = 6
    X_init_a = train_a_X[:n_init_a]
    X_init_b = train_b_X[:n_init_b]
    X_init = np.concatenate((X_init_a, X_init_b), axis=0)
    y_init = np.concatenate((np.repeat(1, n_init_a), np.repeat(0, n_init_b)), axis=0)

    # remaining "unlabeled" instances
    n_a = len(train_a_X)
    n_b = len(train_b_X)
    X_rest = np.concatenate((train_a_X[n_init_a:], train_b_X[n_init_b:]), axis=0)
    y_rest = np.concatenate((np.repeat(1, n_a - n_init_a), np.repeat(0, n_b - n_init_b)), axis=0)

    # train an initial model, used for selecting top n least confident instances
    clf_model_init = LogisticRegression()
    clf_model_init.fit(X_init, y_init)

    # make predictions on the rest of the dataset
    pred_probs = clf_model_init.predict_proba(X_rest)[:, 0]

    # active learning
    accs_active = []
    j = 0
    X_combined_active = X_init.copy()
    y_combined_active = y_init.copy()
    X_rest_active = X_rest.copy()
    y_rest_active = y_rest.copy()
    while j < 20:
        # pick the top n least confident instances and labeled them
        least_conf_idx, rest_idx = get_least_confident_idx(pred_probs, n=5)
        X_new_active = X_rest_active[least_conf_idx]
        y_new_active = y_rest_active[least_conf_idx]

        # update the remaining data
        X_rest_active = X_rest_active[rest_idx]
        y_rest_active = y_rest_active[rest_idx]

        # add new labeled data to existing training set
        X_combined_active = np.concatenate((X_combined_active, X_new_active), axis=0)
        y_combined_active = np.concatenate((y_combined_active, y_new_active), axis=0)
        
        # train a new model
        clf_model_active = LogisticRegression()
        clf_model_active.fit(X_combined_active, y_combined_active)    

        # predict labels on held-out set and compute accuracy
        pred_labels_active = clf_model_active.predict(test_X)
        # accs_active.append(accuracy_score(test_y, pred_labels_active))
        accs_active.append(f1_score(test_y, pred_labels_active))

        # predict class probabilities for the remaining data
        # used for selecting the top n least confident instances in the next iteration
        pred_probs = clf_model_active.predict_proba(X_rest_active)[:, 0]

        j += 1
        
    all_accs_active.append(accs_active)
    
    # traditional learning
    accs_random = []
    k = 0
    n = 5
    X_combined_random = X_init.copy()
    y_combined_random = y_init.copy()
    X_rest_random = X_rest.copy()
    y_rest_random = y_rest.copy()
    while k < 20:
        # randomly pick n instances
        shuffled_indices = np.random.permutation(len(X_rest_random))
        X_new_random = X_rest_random[shuffled_indices[:n]]
        y_new_random = y_rest_random[shuffled_indices[:n]]
        
        # update the remaining data
        X_rest_random = X_rest_random[shuffled_indices[n:]]
        y_rest_random = y_rest_random[shuffled_indices[n:]]
        
        # add new data to existing training set
        X_combined_random = np.concatenate((X_combined_random, X_new_random), axis=0)
        y_combined_random = np.concatenate((y_combined_random, y_new_random), axis=0)

        # train a new model
        clf_model_random = LogisticRegression()
        clf_model_random.fit(X_combined_random, y_combined_random)
        
        # predict labels on held-out set and compute accuracy
        pred_labels_random = clf_model_random.predict(test_X)
        accs_random.append(f1_score(test_y, pred_labels_random))
        # accs_random.append(accuracy_score(test_y, pred_labels_random))

        k += 1
        
    all_accs_random.append(accs_random)
    
    i += 1

# %% [code]
import numpy as np
import pandas as pd
import scipy.stats as st
import plotly.express as px

def compute_ci(data, level=0.05):
    """
    Computes confidence intervals around each mean accuracy.
    """
    ci = []
    for i in range(data.shape[1]):
        ci.append(
            st.t.interval(1 - level, data.shape[0] - 1, loc=np.mean(data[:, i]), scale=st.sem(data[:, i]))
        )
    return np.array(ci)

random_arr = np.array([
    np.arange(5, 105, 5) , 
    np.mean(all_accs_random, axis=0), 
    np.std(all_accs_random, axis=0),
    compute_ci(np.array(all_accs_random))[:, 0],
    compute_ci(np.array(all_accs_random))[:, 1]
])
active_arr = np.array([
    np.arange(5, 105, 5) , 
    np.mean(all_accs_active, axis=0), 
    np.std(all_accs_active, axis=0),
    compute_ci(np.array(all_accs_active))[:, 0],
    compute_ci(np.array(all_accs_active))[:, 1]    
])
arr_combined = np.concatenate((random_arr.T, active_arr.T), axis=0)

# make a pandas DataFrame for plotting
df = pd.DataFrame(arr_combined, columns=['n_train', 'mean_accuracy', 'std', 'ci_low', 'ci_high'])
df['e_minus'] = df['mean_accuracy'] - df['ci_low']
df['e'] = df['ci_high'] - df['mean_accuracy']

df.loc[:, 'strategy'] = 'random'
df.loc[20:, 'strategy'] = 'active'

fig = px.scatter(
    df, x='n_train', y='mean_accuracy',
    error_y_minus='e_minus',
    error_y='e',
    color='strategy',
    color_discrete_sequence=['#0172B2', '#D55E00'],
    labels={
        'n_train': 'Number of labeled instances added',
        'mean_accuracy': 'Mean F1 score on held-out set'
    },
    range_y=[0.75, 0.90]
)
fig.update_traces(
    marker=dict(
        size=12,
            line=dict(
            width=2,
            color='DarkSlateGrey'
        )
    ),
    selector=dict(mode='markers')
)
fig.update_layout(
    font_family='Arial',
    font_size=18,
    showlegend=False,
    xaxis=dict(
        tickmode='linear',
        tick0=10,
        dtick=10,
        ticks='outside'
    ),
    yaxis=dict(
        tickmode='linear',
        tick0=0.55,
        dtick=0.05,
        ticks='outside',
        tickformat='.2f'
    ),
)
fig.update_layout({
    'plot_bgcolor': 'rgba(0, 0, 0, 0)',
})

fig.show()

# %% [code]
```
