## Supplementary material for "An active learning framework improves tumor variant interpretation": Zipped file containing three Jupyter Notebooks with the code used in the study and five source data files required to run the Jupyter Notebooks: active_learning_xpa.ipynb

```
# %% [code]
%reset
import numpy as np
from sklearn.metrics import f1_score, accuracy_score
from sklearn.linear_model import LogisticRegression
import os
import numpy as np
import pandas as pd
import matplotlib.pyplot as plt

# data directory
data_dir = '/path/to/where/xpa_pca_features.csv/is/located/'

# load data file
feature_file = os.path.join(data_dir, 'xpa_pca_features.csv')
xpa_df = pd.read_csv(feature_file)

xpa_train_X = xpa_df[xpa_df['label'] != 'Unknown'].iloc[:, 1:4]
xpa_train_y = xpa_df[xpa_df['label'] != 'Unknown']['label'].apply(lambda x: 1 if x == 'Yes' else 0)
xpa_unknown_X = xpa_df[xpa_df['label'] == 'Unknown'].iloc[:, 1:4]
xpa_unknown_ids = xpa_df[xpa_df['label'] == 'Unknown'].iloc[:, 0]

# logistic regresion model
clf_model = LogisticRegression()
clf_model.fit(xpa_train_X, xpa_train_y)    

# predict labels on test set
pred_probs = clf_model.predict_proba(xpa_unknown_X)

# active learning
def get_least_confident_idx(pred_prob, n=10):
    idx_sorted = np.argsort(np.abs(pred_prob - 0.5))
    return idx_sorted[:n]

least_conf_idx = get_least_confident_idx(pred_probs[:, 0])

# %% [code]
# get the variants that the initial model is most uncertain about 
np.array(xpa_unknown_ids)[least_conf_idx]

# %% [code]
# variants tested by FM-HCR assays and their labels
fm_hcr_variants = [
    'D5Y', 'G6R', 'A18S', 'R30W', 'A60T', 'D70H', 'G72E', 'G73E', 'P94L', 
    'E106G', 'K110E', 'E111A', 'F112C', 'M113I', 'D114Y', 'T125A', 'C126W', 'C126Y', 
    'R130I', 'L138R', 'Y148D', 'D154A', 'F164C', 'V234M', 'H242L', 'R258C', 'K272N'
]
fm_hcr_labels = np.array(
    [
        0, 0, 0, 0, 0, 0, 0, 1, 0, 
        0, 0, 0, 0, 1, 1, 0, 1, 1, 
        0, 1, 1, 0, 0, 0, 0, 0, 0
    ]
)

# %% [markdown]
### Active learning

# %% [code]
# start fresh with the original data set
feature_file = os.path.join(data_dir, 'xpa_pca_features.csv')
xpa_df = pd.read_csv(feature_file)

# label the variants the inital model was most uncertain about
for x in ['H242L', 'D70H', 'E111A', 'D154A', 'T125A', 'E106G']:
    xpa_df.loc[xpa_df['variant'] == x, 'label'] = 'No'
for y in ['L138R']:
    xpa_df.loc[xpa_df['variant'] == y, 'label'] = 'Yes'

# add these variants to the training set
xpa_train_X = xpa_df[xpa_df['label'] != 'Unknown'].iloc[:, 1:4]
xpa_train_y = xpa_df[xpa_df['label'] != 'Unknown']['label'].apply(lambda x: 1 if x == 'Yes' else 0)
xpa_unknown_X = xpa_df[xpa_df['label'] == 'Unknown'].iloc[:, 1:4]

# train an updated logistic regresion model
clf_model_active = LogisticRegression()
clf_model_active.fit(xpa_train_X, xpa_train_y)

# %% [markdown]
### Compare active learning with "random" learning

# %% [code]
# set random seed to reproduce results
np.random.seed(0)

# start fresh with the original data set
feature_file = os.path.join(data_dir, 'xpa_pca_features.csv')
xpa_df = pd.read_csv(feature_file)

active_f1_scores = []
random_f1_scores = []
k = 0
while k < 100:
    idx = np.random.choice(len(fm_hcr_variants), 7, replace=False)
    random_variants = [fm_hcr_variants[i] for i in idx]
    random_labels = [fm_hcr_labels[i] for i in idx]

    for x, y in zip(random_variants, random_labels):
        z = 'Yes' if y == 1 else 'No'
        xpa_df.loc[xpa_df['variant'] == x, 'label'] = z

    xpa_train_X = xpa_df[xpa_df['label'] != 'Unknown'].iloc[:, 1:4]
    xpa_train_y = xpa_df[xpa_df['label'] != 'Unknown']['label'].apply(lambda x: 1 if x == 'Yes' else 0)
    xpa_unknown_X = xpa_df[xpa_df['label'] == 'Unknown'].iloc[:, 1:4]
    xpa_unknown_ids = xpa_df[xpa_df['label'] == 'Unknown'].iloc[:, 0]

    # logistic regresion model
    clf_model_random = LogisticRegression()
    clf_model_random.fit(xpa_train_X, xpa_train_y)
    
    # create a test set
    test_set_variants = set(fm_hcr_variants) - set(
        ['H242L', 'D70H', 'E111A', 'D154A', 'T125A', 'E106G', 'L138R'] + random_variants
    )
    test_set_idx = [fm_hcr_variants.index(x) for x in test_set_variants]
    test_set_y = [fm_hcr_labels[i] for i in test_set_idx]
    test_set_X = xpa_df.iloc[
        [list(xpa_df['variant']).index(x) for x in test_set_variants], 
        1:4
    ]
    
    # evaluate actively learned model and "random" model
    active_f1_scores.append(
        f1_score(clf_model_active.predict(test_set_X), test_set_y)
    )
    random_f1_scores.append(
        f1_score(clf_model_random.predict(test_set_X), test_set_y)
    )
    
    k += 1

# %% [code]
np.mean(active_f1_scores)

# %% [code]
np.mean(random_f1_scores)
```
