## Supplementary Figures S1-S4 and Supplementary Tables S1-S8 for "An active learning framework improves tumor variant interpretation"

### Contents

Supplementary Figure S1. Spearman's rank correlation of input features.
Supplementary Figure S2. Principal component analyses of XPA variant input features.
Supplementary Figure S3. Active versus traditional learning using additional synthetic and deep mutational scanning datasets with increasing amounts of labeled training instances.
Supplementary Figure S4. Semi-supervised label spreading algorithm approach to predict NER-deficient variants.
Supplementary Table S1. Deep mutational scanning data summary.
Supplementary Table S2. Labeled XPA variant sources, reported phenotypes, and references.
Supplementary Table S3. Unlabeled XPA variant sources, reported phenotypes, and references.
Supplementary Table S4. dbNSFP data for labeled XPA variants.
Supplementary Table S5. Algorithm predictions for XPA VUS.
Supplementary Table S6. Algorithm predictions for labeled XPA variants.
Supplementary Table S7. Average FM-HCR relative reporter expression normalized to wild-type compared with dbNSFP data for selected XPA variants.
Supplementary Table S8. Average FM-HCR relative reporter expression normalized to wild-type compared with logistic regression model predictions for selected XPA VUS.

### Additional Files

All code and required source data are available as a zipped folder "jupyter_notebooks_and_source_data.zip" with the following contents:

- Code provided as a Jupyter Notebook to run the active learning simulation for DMS datasets: active_learning_dms.ipynb
- Accompanying source data (input features) for each of the four DMS proteins: cyp2c9_pca_features.csv; nudt15_pca_features.csv; pten_pca_features.csv; tpmt_pca_features.csv
- Code provided as a Jupyter Notebook to generate the synthetic datasets and run the active learning simulation: active_learning_synthetic_data.ipynb
- Code provided as a Jupyter Notebook to run the active learning simulation for the XPA dataset: active_learning_xpa.ipynb
- Accompanying source data (input features) for XPA: xpa_pca_features.csv

**Supplementary Figures**

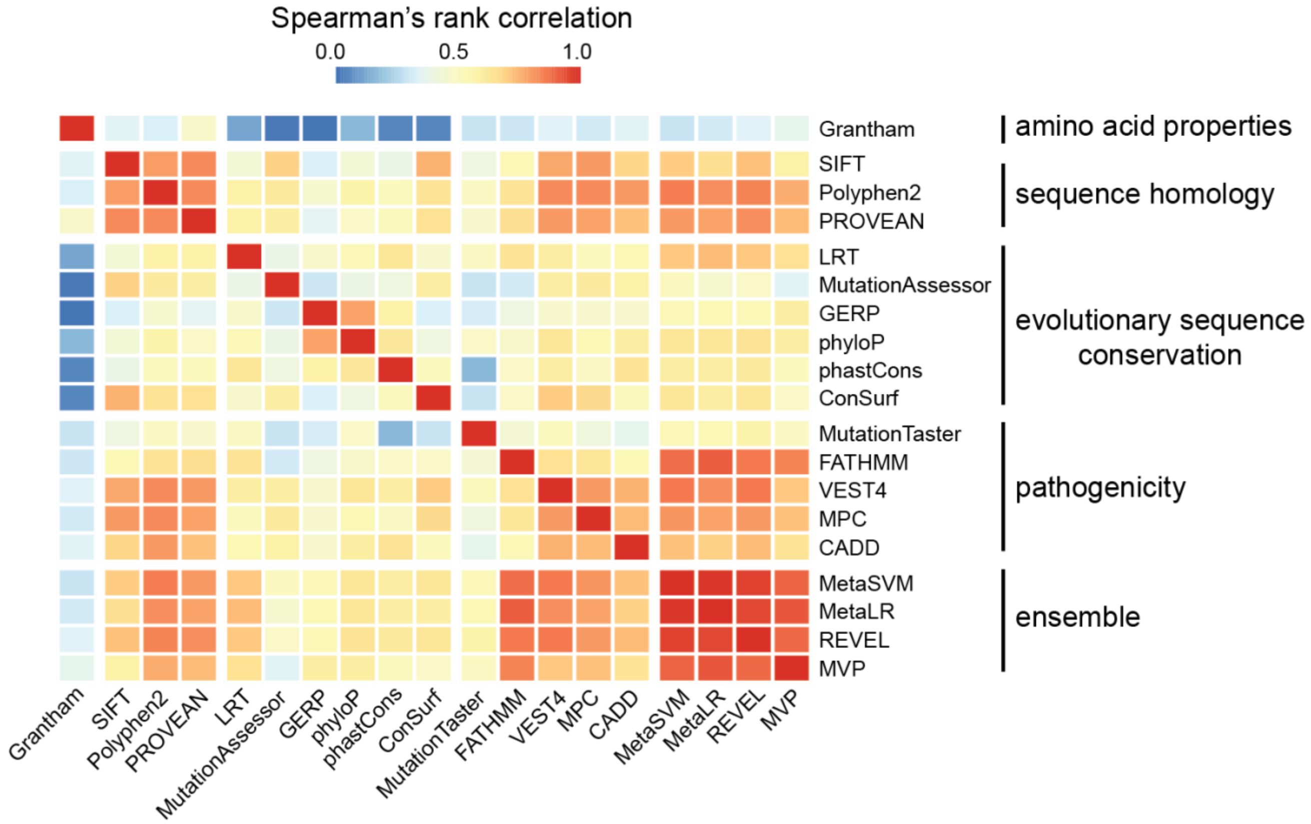

**Supplementary Figure S1. Spearman's rank correlation of input features.**
Heatmap of pairwise correlations of all 19 features for each XPA variant.

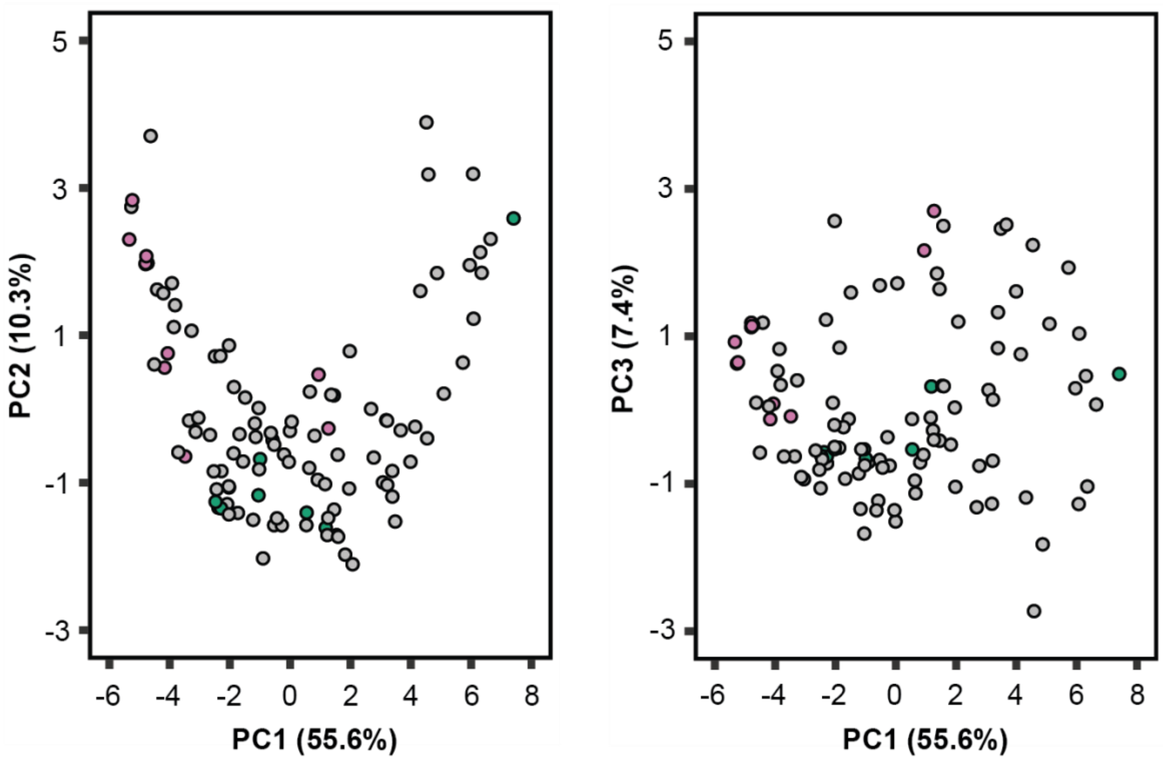

**Supplementary Figure S2. Principal component analyses of XPA variant input features.**
Principal component analysis plots showing principal component (PC) 1 versus PC2 (left) and PC1 versus PC3 (right) for the input features of all XPA variants. Together, the first three PCs describe 73.3% of the variance in the dataset. Labeled NER-deficient variants shown in pink, labeled NER-proficient variants shown in green, unlabeled variants shown in grey.

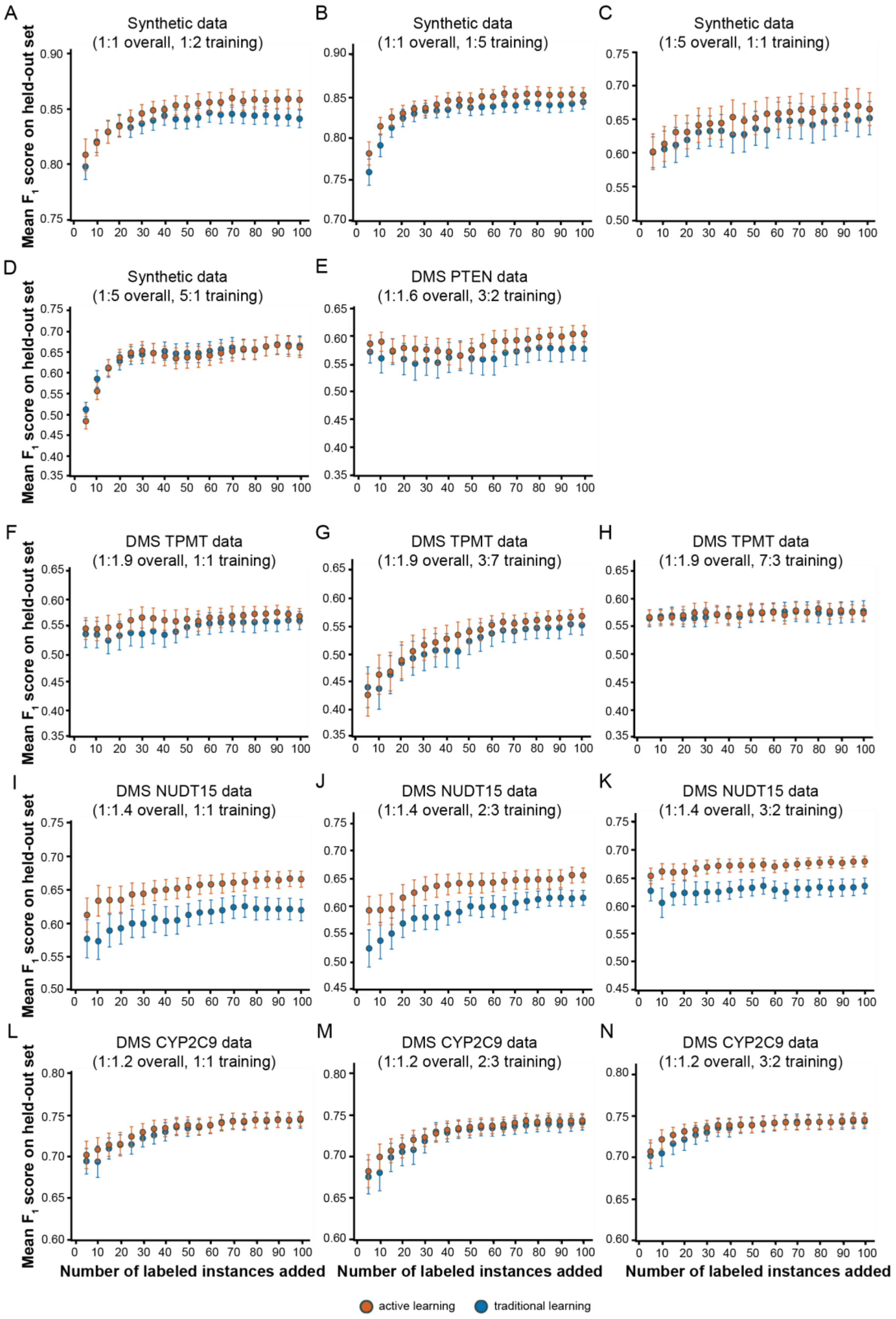

**Supplementary Figure S3. Active versus traditional learning using additional synthetic and deep mutational scanning datasets with increasing amounts of labeled training instances.**

Mean $F_1$ evaluation score comparing active and traditional learning for: synthetic datasets with balanced class ratios (1:1) in the overall data and skewed class ratios in the initial labeled training set (1:2) in *A* and (1:5) in *B*, or skewed class ratios (1:5) in the overall data and a balanced initial labeled training set (1:1) in *C* and a skewed class ratio in the initial labeled training set (5:1) in *D*; a DMS PTEN dataset with a skewed class ratio (3:2) in the initial labeled training set in *E*; a DMS TPMT dataset with a balanced initial labeled training set (1:1) in *F* and skewed class ratios in the initial labeled training set (3:7) in *G* and (7:3) in *H*; a DMS NUDT15 dataset with a balanced initial labeled training set (1:1) in *I* and skewed class ratios in the initial labeled training set (2:3) in *J* and (3:2) in *K*; and a DMS CYP2CP9 dataset with a balanced initial labeled training set (1:1) in *L* and skewed class ratios in the initial labeled training set (2:3) in *M* and (3:2) in *N*. Error bars indicate 95% confidence intervals around the mean $F_1$ score. All initial labeled pools had ten instances to start except for the skewed synthetic dataset in *A*-*D*, which had 12 instances to maintain the 1:5 ratio with sufficient starting numbers of instances in both classes. See **Supplementary Table S1** for additional details regarding the composition of the DMS datasets.

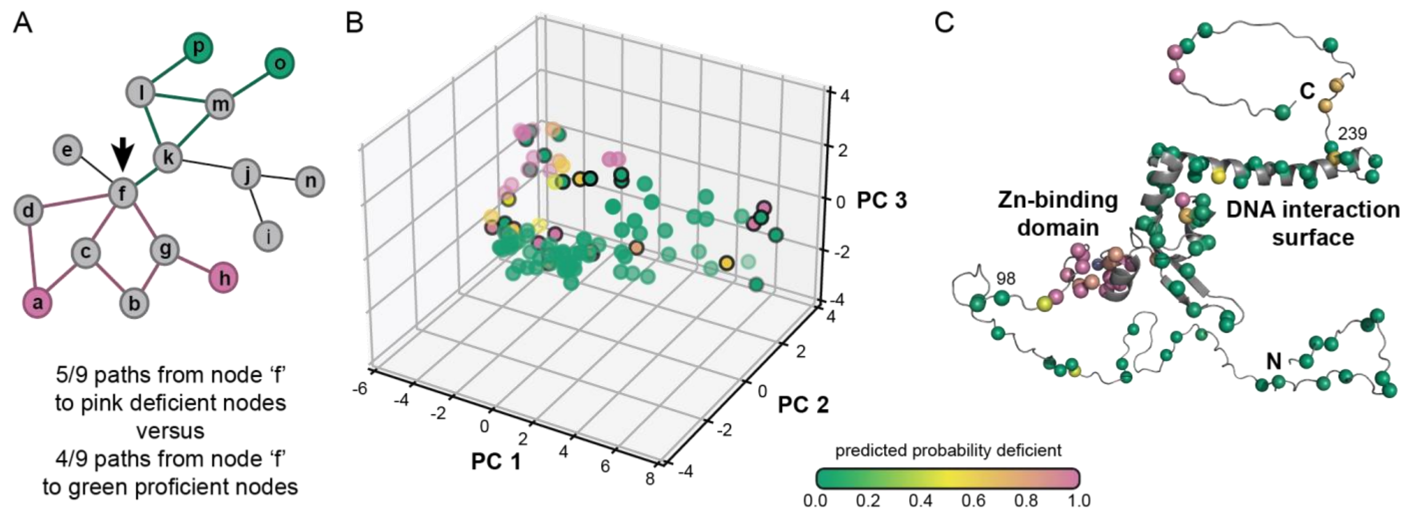

**Supplementary Figure S4. Semi-supervised label spreading algorithm approach to predict NER-deficient variants.**

*A*, Schematic representation of the label spreading algorithm. Here, the probability of node F being NER-deficient is predicted as the fraction of paths between node F and known NER-deficient variants (pink) versus the fraction of paths between node F and known NER-proficient nodes (green). *B*, Effects of XPA variants on NER activity predicted by the label spreading algorithm. Input features are the first three principal components from a principal component analysis (PCA) of the original set of 19 features from dbNSFP. Variants selected for functional validation by FM-HCR outlined in black: D5Y, G6R, A18S, R30W, A60T, D70H, G72E, G73E, P94L, E106G, K110E, E111A, F112C, M113I, D114Y, T125A, C126W, C126Y, R130I, L138R, Y148D, D154A, F164C, V234M, H242L, R258C, and K272N. *C*, Representative model of full-length XPA with variants of interest depicted as spheres and colored according to scheme in *B*. The precise fold and orientation of the flexible N- and C-termini regions are not known and are shown only for representative purposes.

**Supplementary Tables**

**Supplementary Table S1. Deep mutational scanning data summary.**

| Protein | # Low-Abundance Variants (class 1) | # Wild-type-like Variants (class 2) | Class Ratio | Abundance Score Cutoff | References |
|---|---|---|---|---|---|
| **PTEN** | 522 | 844 | skewed (1:1.6) | 0.71 | PMID: 29785012 |
| **TPMT** | 385 | 746 | skewed (1:1.9) | 0.72 | PMID: 29785012 |
| **NUDT15** | 366 | 524 | skewed (1:1.4) | 0.72 | PMID: 32094176 |
| **CYP2C9** | 659 | 778 | skewed (1:1.2) | 0.77 | bioRxiv doi: 10.1101/2021.03.12.435209 |

**Supplementary Table S2. Labeled XPA variant sources, reported phenotypes, and references.**

| Variant | Variant Source | Training Label | Cell Survival/*in vivo* Repair Defect | *in vitro* Repair Defect | DNA Binding Defect | Protein Interaction Defect | PMIDs | | | | |
|---|---|---|---|---|---|---|---|---|---|---|---|
| C105S | Synthetic | Deficient | Moderate | - | - | - | 8504220 | 1601884 | - | - | - |
| E106A | Synthetic | Deficient | - | None | None | None (RPA70) | 21148310 | - | - | - | - |
| E106K | Synthetic | Deficient | Moderate | None | - | Modest (RPA70AB) | 31925419 | - | - | - | - |
| C108F | XP disease | Deficient | Severe | - | - | - | 1339397 | 8504220 | 24704021 | 25326635 | - |
| C108G | Synthetic | Deficient | Moderate | - | - | - | 1601884 | - | - | - | - |
| C108S | Synthetic | Deficient | Moderate | - | - | - | 8504220 | 1601884 | - | - | - |
| C126S | Synthetic | Deficient | Moderate | - | - | - | 8504220 | 1601884 | - | - | - |
| C129S | Synthetic | Deficient | Moderate | - | - | - | 8504220 | 1601884 | - | - | - |
| K141E | Synthetic | Proficient | None | - | Severe | Modest (ERCC1-XPF) | 16491090 | - | - | - | - |
| K145E | Synthetic | Proficient | None | - | Severe | - | 16491090 | - | - | - | - |
| K151E | Synthetic | Proficient | None | - | - | - | 16491090 | - | - | - | - |
| C153S | Synthetic | Proficient | None | - | - | - | 8504220 | 1601884 | - | - | - |
| K179E | Synthetic | Proficient | None | - | Severe | Modest (ERCC1-XPF) | 16491090 | - | - | - | - |
| K204E | Synthetic | Proficient | None | - | - | - | 16491090 | - | - | - | - |
| R228Q | Healthy | Proficient | None | - | - | - | 15661657 | 12509227 | 16491090 | 10862089 | 24728327 |
| V234L | Healthy | Proficient | None | - | - | - | 15661657 | 12509227 | - | - | - |
| H244R | XP disease | Deficient | Moderate | - | - | - | 8504220 | 1372103 | 9753735 | - | - |
| C261S | Synthetic | Deficient | Moderate | - | - | - | 8504220 | 1601884 | - | - | - |
| C264S | Synthetic | Deficient | Moderate | - | - | - | 8504220 | 1601884 | 9753735 | - | - |

**Supplementary Table S3. Unlabeled XPA variant sources, reported phenotypes, and references.**

| Variant | Variant Source | Cancer Type (for Tumor Variants) | Frequency (for Tumor Variants) | Cell Survival/*in vivo* Repair Defect | *in vitro* Repair Defect | DNA Binding Defect | Protein Interaction Defect | PMIDs | | | |
|---|---|---|---|---|---|---|---|---|---|---|---|
| **A3G** | CBioPortal, ICGC | BLCA | 1 | - | - | - | - | - | - | - | - |
| **D5Y** | CBioPortal, COSMIC | COAD | 2 | - | - | - | - | - | - | - | - |
| | GENIE | Serous Ovarian Cancer | 2 | - | - | - | - | - | - | - | - |
| **G6R** | PanCanAtlas, CBioPortal, COSMIC | LUSC | 1 | - | - | - | - | - | - | - | - |
| **G6W** | PanCanAtlas, CBioPortal, COSMIC, ICGC | LUSC | 1 | - | - | - | - | - | - | - | - |
| **E10K** | PanCanAtlas, CBioPortal, COSMIC, ICGC | BLCA | 1 | - | - | - | - | - | - | - | - |
| **A12V** | CBioPortal, COSMIC | COAD | 1 | - | - | - | - | - | - | - | - |
| **P17H** | PanCanAtlas, CBioPortal, COSMIC, ICGC | PRAD | 1 | - | - | - | - | - | - | - | - |
| **A18S** | COSMIC, ICGC | THCA | 1 | - | - | - | - | - | - | - | - |
| **S23L** | GENIE | LUSC | 1 | - | - | - | - | - | - | - | - |
| | PanCanAtlas, CBioPortal, ICGC | HNSC | 1 | - | - | - | - | - | - | - | - |
| **R25W** | CBioPortal, DepMap/CCLE | COAD | 1 | - | - | - | - | - | - | - | - |
| **R30W** | COSMIC | THCA | 1 | - | - | - | - | - | - | - | - |
| **Q33L** | COSMIC | COAD | 1 | - | - | - | - | - | - | - | - |
| **L43P** | CBioPortal | SARC | 1 | - | - | - | - | - | - | - | - |
| **R46W** | COSMIC, ICGC | PRAD | 1 | - | - | - | - | - | - | - | - |
| **Y48F** | CBioPortal, DepMap/CCLE | HNSC | 1 | - | - | - | - | - | - | - | - |
| **A60D** | PanCanAtlas, CBioPortal, COSMIC, ICGC | STAD | 1 | - | - | - | - | - | - | - | - |
| **A60T** | COSMIC, DepMap/CCLE, CBioPortal | COAD | 1 | - | - | - | - | - | - | - | - |

| | | | | | | | | | | | |
|---|---|---|---|---|---|---|---|---|---|---|---|
| **A64T** | PanCanAtlas, CBioPortal, COSMIC | STAD | 2 | - | - | - | - | - | - | - | - |
| | COSMIC | Skin | 2 | - | - | - | - | - | - | - | - |
| **A65T** | PanCanAtlas, CBioPortal, COSMIC, ICGC | COAD | 1 | - | - | - | - | - | - | - | - |
| **A65V** | COSMIC | COAD | 1 | - | - | - | - | - | - | - | - |
| **D70H** | DepMap/CCLE, CBioPortal | Multiple Myeloma | 1 | - | - | - | - | - | - | - | - |
| **D70N** | CBioPortal, DepMap/CCLE | Unknown (BLCA) | 1 | - | - | - | - | - | - | - | - |
| **T71K** | COSMIC, ICGC | Liver | 1 | - | - | - | - | - | - | - | - |
| **G72E** | COSMIC, ICGC | BRCA | 1 | - | - | - | - | - | - | - | - |
| **G73E** | PanCanAtlas, CBioPortal, COSMIC, ICGC | LUAD | 1 | - | - | - | - | - | - | - | - |
| **E81D** | PanCanAtlas, CBioPortal | UCEC | 1 | - | - | - | - | - | - | - | - |
| **P94L** | XP disease | - | - | - | - | - | - | 8504220 | - | - | - |
| **P96L** | COSMIC | SKCM | 1 | - | - | - | - | - | - | - | - |
| **P96S** | COSMIC | SKCM | 1 | - | - | - | - | - | - | - | - |
| **M98I** | CBioPortal, ICGC | BLCA | 1 | - | - | - | - | - | - | - | - |
| **D101A** | Synthetic | - | - | - | None | None | None (RPA70) | 21148310 | - | - | - |
| **Y102C** | CBioPortal, COSMIC | COAD | 1 | - | - | - | - | - | - | - | - |
| **Y102H** | COSMIC | GBM | 1 | - | - | - | - | - | - | - | - |
| **E106G** | COSMIC | COAD | 1 | - | - | - | - | - | - | - | - |
| **K110E** | PanCanAtlas, CBioPortal | UCEC | 1 | - | - | - | - | - | - | - | - |
| **E111A** | Synthetic | - | - | - | None | None | None (RPA70) | 21148310 | - | - | - |
| **F112C** | PanCanAtlas | UCEC | 1 | - | - | - | - | - | - | - | - |
| **M113I** | GENIE | Squamous Cell Carcinoma | 2 | - | - | - | - | - | - | - | - |
| | CBioPortal, ICGC | SKCM | 2 | - | - | - | - | - | - | - | - |
| **D114A** | Synthetic | - | - | - | Severe | None | None (RPA70) | 21148310 | - | - | - |
| **D114Y** | COSMIC, ICGC | THCA | 1 | - | | | | - | - | - | - |

| | | | | | | | | | | | |
|---|---|---|---|---|---|---|---|---|---|---|---|
| **D122A** | Synthetic | - | - | - | None | None | None (RPA70) | 21148310 | - | - | - |
| **T125A** | PanCanAtlas, CBioPortal, ICGC | UCEC | 1 | - | - | - | - | - | - | - | - |
| **C126W** | GENIE | Medulloblastoma | 1 | - | - | - | - | - | - | - | - |
| **C126Y** | XP disease | - | - | - | - | - | - | 9671271 | - | - | - |
| **D127A** | Synthetic | - | - | - | None | None | None (RPA70) | 21148310 | - | - | - |
| **R130I** | CBioPortal, ICGC | SKCM | 1 | - | - | - | - | - | - | - | - |
| **R130K** | XP disease | - | - | - | - | - | - | 8504220 | 9671271 | | |
| **L138R** | PanCanAtlas, CBioPortal, ICGC | STAD | 1 | - | - | - | - | - | - | - | - |
| **T142A** | Synthetic | - | - | - | Severe | None | Severe (RPA70) | 21148310 | - | - | - |
| **Q146K** | CBioPortal, COSMIC | BRCA | 1 | - | | | | - | - | - | - |
| **Y148D** | CBioPortal | SARC | 1 | - | | | | - | - | - | - |
| **D154A** | Synthetic | - | - | - | None | None | None (RPA70) | 21148310 | - | - | - |
| **E159A** | CBioPortal, COSMIC, ICGC | BRCA | 1 | - | - | - | - | - | - | - | - |
| **E159G** | DepMap/CCLE, CBioPortal | THCA | 1 | - | - | - | - | - | - | - | - |
| **L162V** | PanCanAtlas, CBioPortal, COSMIC, ICGC | UCEC | 1 | - | - | - | - | - | - | - | - |
| **F164C** | GENIE | COAD | 2 | - | - | - | - | - | - | - | - |
| | CBioPortal, COSMIC | COAD | 2 | - | - | - | - | - | - | - | - |
| **V166A** | CBioPortal | DLBC | 1 | - | - | - | - | - | - | - | - |
| **K168N** | PanCanAtlas | UCEC | 1 | - | - | - | - | - | - | - | - |
| **H172Y** | GENIE | Clear Cell Ovarian Cancer | 2 | - | - | - | - | - | - | - | - |
| | PanCanAtlas, CBioPortal, ICGC | UCEC | 2 | - | - | - | - | - | - | - | - |
| **S173L** | COSMIC, ICGC | ESCA/ESAD | 1 | - | - | - | - | - | - | - | - |
| **Q185H** | XP disease | - | - | - | - | - | - | 25525159 | 8504220 | | |
| **V187A** | PanCanAtlas, CBioPortal, COSMIC, ICGC | STAD | 1 | - | - | - | - | - | - | - | - |
| **V187L** | COSMIC | STAD | 1 | - | - | - | - | - | - | - | - |

| | | | | | | | | | | | |
|---|---|---|---|---|---|---|---|---|---|---|---|
| **S190A** | Healthy | - | - | - | - | - | - | 24728327 | | | |
| **S190F** | PanCanAtlas, CBioPortal, COSMIC, ICGC | CESC | 1 | - | - | - | - | - | - | - | - |
| **L191V** | CBioPortal | Mixed Cancer Types | 1 | - | - | - | - | - | - | - | - |
| **E198A** | Synthetic | - | - | - | None | None | None (RPA70) | 21148310 | | | |
| **A199V** | CBioPortal, COSMIC | COAD | 1 | - | | | | - | - | - | - |
| **A203S** | CBioPortal | PRAD | 1 | - | | | | - | - | - | - |
| **E205A** | Synthetic | - | - | - | None | None | None (RPA70) | 21148310 | - | - | - |
| **R207G** | Synthetic | - | - | Conflicting Data | Severe | None | Severe (DDB1/2) | 8464385 | 19056823 | 16491090 | 9753735 |
| **R207Q** | GENIE | BRCA | 2 | - | - | - | - | - | - | - | - |
| | PanCanAtlas, CBioPortal, COSMIC, ICGC | UCEC | 2 | - | - | - | - | - | - | - | - |
| **R211Q** | GENIE | Uterine Endometrioid Carcinoma | 2 | - | - | - | - | - | - | - | - |
| | CBioPortal, ICGC | COAD | 2 | - | - | - | - | - | - | - | - |
| **E212K** | GENIE | Cutaneous Squamous Cell Carcinoma | 2 | - | - | - | - | - | - | - | - |
| | CBioPortal, COSMIC | LUAD | 2 | - | - | - | - | - | - | - | - |
| **K213Q** | PanCanAtlas, CBioPortal, ICGC | COAD | 1 | - | - | - | - | - | - | - | - |
| **Q216K** | COSMIC | THCA | 1 | - | - | - | - | - | - | - | - |
| **K217N** | PanCanAtlas, CBioPortal, ICGC | UCEC | 2 | - | - | - | - | - | - | - | - |
| | CBioPortal, COSMIC | COAD | 2 | - | - | - | - | - | - | - | - |
| **K221E** | COSMIC | BRCA | 1 | - | - | - | - | - | - | - | - |
| **K222E** | Synthetic | - | - | - | - | Severe | - | 28860187 | 31925419 | - | - |
| **R227W** | GENIE | COAD | 4 | - | - | - | - | - | - | - | - |
| | GENIE | COAD | 4 | - | - | - | - | - | - | - | - |
| | CBioPortal, COSMIC | COAD | 4 | - | - | - | - | - | - | - | - |
| | CBioPortal, COSMIC | COAD | 4 | - | - | - | - | - | - | - | - |

| | | | | | | | | | | | |
|---|---|---|---|---|---|---|---|---|---|---|---|
| **S233N** | CBioPortal, DepMap/CCLE | Mixed Cancer Types | 1 | - | - | - | - | - | - | - | - |
| **V234M** | GENIE | Papillary Thyroid Cancer | 2 | - | - | - | - | - | - | - | - |
| | PanCanAtlas, CBioPortal, COSMIC, ICGC | UCEC | 2 | - | - | - | - | - | - | - | - |
| **T239M** | PanCanAtlas, CBioPortal, COSMIC, ICGC | UCEC | 1 | - | - | - | - | - | - | - | - |
| **H242L** | PanCanAtlas, CBioPortal, COSMIC, ICGC | UCEC | 1 | - | - | - | - | - | - | - | - |
| **E249Q** | GENIE | Ovarian Cancer | 2 | - | - | - | - | - | - | - | - |
| | CBioPortal, COSMIC | PRAD | 2 | - | - | - | - | - | - | - | - |
| **R258C** | GENIE | Myeloid Neoplasm | 3 | - | - | - | - | - | - | - | - |
| | CBioPortal, COSMIC | COAD | 3 | - | - | - | - | - | - | - | - |
| | PanCanAtlas, CBioPortal, ICGC | UCEC | 3 | - | - | - | - | - | - | - | - |
| **R258H** | PanCanAtlas, CBioPortal, ICGC | GBM | 3 | - | - | - | - | - | - | - | - |
| | CBioPortal | B-Lymphoblastic Leukemia/Lymphoma | 3 | - | - | - | - | - | - | - | - |
| | COSMIC, ICGC | COAD | 3 | - | - | - | - | - | - | - | - |
| **T260N** | PanCanAtlas, CBioPortal, ICGC | UCEC | 1 | - | - | - | - | - | - | - | - |
| **K272N** | PanCanAtlas, CBioPortal, ICGC | GBM | 1 | - | - | - | - | - | - | - | - |

**Supplementary Table S4. dbNSFP data for labeled XPA variants.**

| | | Amino Acid Properties | Sequence Homology | | | Evolutionary Sequence Conservation | | | | | |
|---|---|---|---|---|---|---|---|---|---|---|---|
| Variant | Training Label | Grantham | SIFT | Polyphen2 | PROVEAN | LRT | MutationAssessor | GERP | phyloP | phastCons | ConSurf |
| C105S | Deficient | 112 | 0 | 0.999 | -9.72 | 0 | 2.265 | 4.88 | 7.276 | 1 | -1.2 |
| E106A | Deficient | 107 | 0.711 | 0.328 | -2.47 | 0 | 1.445 | 5.25 | 5.491 | 1 | 1.165 |
| E106K | Deficient | 56 | 0.863 | 0.264 | -1.67 | 0 | 1.445 | 5.77 | 7.276 | 1 | 1.165 |
| C108F | Deficient | 205 | 0 | 1 | -10.86 | 0 | 2.265 | 5.25 | 7.096 | 1 | -1.159 |
| C108G | Deficient | 159 | 0 | 0.999 | -11.85 | 0 | 2.265 | 5.25 | 8.38 | 1 | -1.159 |
| C108S | Deficient | 112 | 0 | 0.998 | -9.87 | 0 | 2.265 | 5.25 | 7.096 | 1 | -1.159 |
| C126S | Deficient | 112 | 0 | 0.999 | -9.89 | 0 | 2.265 | 5.25 | 6.874 | 1 | -1.202 |
| C129S | Deficient | 112 | 0.003 | 0.988 | -9.76 | 0 | 2.25 | 5.25 | 6.874 | 1 | -1.201 |
| K141E | Proficient | 56 | 0.006 | 0.778 | -3.07 | 1.8E-05 | 2.515 | 5.34 | 4.721 | 1 | -0.6953 |
| K145E | Proficient | 56 | 0.003 | 0.897 | -3.83 | 0 | 3.24 | 5.34 | 7.628 | 1 | -1.051 |
| K151E | Proficient | 56 | 0.002 | 0.988 | -3.69 | 0 | 2.955 | 5.34 | 7.628 | 1 | -0.9953 |
| C153S | Proficient | 112 | 0.002 | 0.794 | -8.53 | 4.6E-05 | 1.935 | 5.34 | 5.727 | 1 | -0.3354 |
| K179E | Proficient | 56 | 0 | 0.909 | -4 | 2E-06 | 3.02 | 5.63 | 7.628 | 1 | -0.9355 |
| K204E | Proficient | 56 | 0.077 | 0.486 | -2.53 | 5E-06 | 2.73 | 4.98 | 4.06 | 1 | -0.01684 |
| R228Q | Proficient | 43 | 0.324 | 0.749 | -0.34 | 4.1E-05 | 1.445 | 5.32 | 5.303 | 1 | -0.557 |
| V234L | Proficient | 32 | 1 | 0.004 | 1.01 | 0.067075 | -0.64 | 0.0447 | -0.24 | 0.009 | -0.2359 |
| H244R | Deficient | 29 | 0 | 0.997 | -7.37 | 3E-06 | 3.31 | 5.32 | 7.072 | 1 | -1.18 |
| C261S | Deficient | 112 | 0 | 0.996 | -8.96 | 1E-06 | 2.77 | 5.32 | 6.976 | 1 | -1.159 |
| C264S | Deficient | 112 | 0 | 0.996 | -8.59 | 0 | 3.32 | 5.32 | 6.976 | 1 | -1.153 |

| | | Pathogenicity | | | | | Ensemble | | | |
|---|---|---|---|---|---|---|---|---|---|---|
| Variant | Training Label | MutationTaster | FATHMM | VEST4 | MPC | CADD | MetaSVM | MetaLR | REVEL | MVP |
| C105S | Deficient | 1 | -5.5 | 0.991 | 1.047324358 | 3.775677 | 1.0872 | 0.9742 | 0.971 | 0.975240068 |
| E106A | Deficient | 1 | -1.84 | 0.349 | 0.377018116 | 3.247832 | -0.1912 | 0.5552 | 0.462 | 0.921291597 |
| E106K | Deficient | 1 | -1.82 | 0.354 | 0.404426802 | 3.055115 | -0.272 | 0.4453 | 0.439 | 0.921852769 |
| C108F | Deficient | 1 | -5.51 | 0.968 | 1.177491728 | 3.967267 | 1.0856 | 0.9766 | 0.924 | 0.997153832 |
| C108G | Deficient | 1 | -5.5 | 0.949 | 1.053085251 | 4.328254 | 1.0852 | 0.9746 | 0.979 | 0.99700408 |
| C108S | Deficient | 1 | -5.5 | 0.96 | 1.047324358 | 3.850675 | 1.0856 | 0.9766 | 0.927 | 0.998154988 |
| C126S | Deficient | 1 | -5.5 | 0.982 | 1.047324358 | 4.003157 | 1.0856 | 0.9766 | 0.923 | 0.966221379 |
| C129S | Deficient | 1 | -5.5 | 0.978 | 1.040315251 | 3.901357 | 1.0862 | 0.9763 | 0.911 | 0.970443012 |
| K141E | Proficient | 0.993211 | -0.14 | 0.628 | 1.075489361 | 4.192903 | -0.0991 | 0.4049 | 0.346 | 0.850903602 |
| K145E | Proficient | 1 | -0.16 | 0.838 | 1.070717868 | 4.289727 | 0.1149 | 0.4752 | 0.739 | 0.879448877 |
| K151E | Proficient | 1 | 0.04 | 0.776 | 1.070717868 | 4.463131 | 0.0512 | 0.4699 | 0.625 | 0.88877857 |

| | | | | | | | | | | |
|---|---|---|---|---|---|---|---|---|---|---|
| **C153S** | Proficient | 1 | 0.27 | 0.794 | 0.447225067 | 4.102858 | -0.4628 | 0.3017 | 0.41 | 0.900229055 |
| **K179E** | Proficient | 1 | 0.03 | 0.838 | 1.103808958 | 4.198148 | 0.0516 | 0.4615 | 0.648 | 0.919252916 |
| **K204E** | Proficient | 1 | 0.11 | 0.555 | 0.746878719 | 3.452975 | -0.5422 | 0.2463 | 0.172 | 0.83223147 |
| **R228Q** | Proficient | 0.999976 | 0.19 | 0.571 | 0.348634747 | 3.957845 | -0.8361 | 0.2369 | 0.164 | 0.778699478 |
| **V234L** | Proficient | 1 | 0.3 | 0.022 | 0.261781093 | 0.691653 | -1.0161 | 0.0391 | 0.042 | 0.266385637 |
| **H244R** | Deficient | 1 | -1.16 | 0.924 | 1.043197875 | 3.903625 | 0.5122 | 0.6735 | 0.87 | 0.978861589 |
| **C261S** | Deficient | 1 | -2.29 | 0.959 | 1.047324358 | 3.876653 | 0.8091 | 0.8267 | 0.78 | 0.996031426 |
| **C264S** | Deficient | 1 | -2.26 | 0.961 | 1.047324358 | 3.930123 | 0.7878 | 0.8206 | 0.814 | 0.988095081 |

Where possible, cells are colored to reflect VEP tool predictions (pink for deleterious variants and green for tolerated variants).

**Supplementary Table S5. Algorithm predictions for XPA VUS.**

| Variant | Label Spreading Prediction | Label Spreading P(NER-deficient) | Logistic Regression Class Probability (NER-Deficient) |
|---|---|---|---|
| A3G | Proficient | 0 | 0.007 |
| D5Y | Proficient | 0 | 0.002 |
| G6R | Proficient | 0 | 0.022 |
| G6W | Proficient | 0 | 0.009 |
| E10K | Proficient | 0 | 0.065 |
| A12V | Proficient | 0 | 0.058 |
| P17H | Proficient | 0 | 0.015 |
| A18S | Proficient | 0 | 0.066 |
| S23L | Proficient | 0 | 0.01 |
| R25W | Proficient | 3.63398E-07 | 0.109 |
| R30W | Proficient | 9.74785E-06 | 0.098 |
| Q33L | Proficient | 0 | 0.031 |
| L43P | Proficient | 0.05084453 | 0.229 |
| R46W | Proficient | 0 | 0.03 |
| Y48F | Proficient | 0 | 0.06 |
| A60D | Proficient | 0 | 0.018 |
| A60T | Proficient | 0 | 0.038 |
| A64T | Proficient | 0 | 0.019 |
| A65T | Proficient | 0 | 0.048 |
| A65V | Proficient | 0 | 0.05 |
| D70H | Proficient | 0.4311164 | 0.418 |
| D70N | Proficient | 2.65583E-07 | 0.284 |
| T71K | Proficient | 0 | 0.041 |
| G72E | Proficient | 0 | 0.06 |
| G73E | Proficient | 0.000369117 | 0.186 |
| E81D | Proficient | 0 | 0.155 |
| P94L | Proficient | 9.74785E-06 | 0.208 |
| P96L | Proficient | 0.000335712 | 0.212 |
| P96S | Proficient | 2.80678E-07 | 0.158 |
| M98I | Proficient | 0 | 0.4 |
| D101A | Proficient | 0.4220661 | 0.829 |
| Y102C | Deficient | 0.9903235 | 0.932 |
| Y102H | Deficient | 0.6989253 | 0.899 |
| E106G | Deficient | 0.9939312 | 0.663 |
| K110E | Deficient | 0.9572696 | 0.738 |
| E111A | Deficient | 0.8518371 | 0.583 |
| F112C | Deficient | 0.9996084 | 0.962 |
| M113I | Deficient | 0.8548851 | 0.75 |
| D114A | Deficient | 0.9998345 | 0.974 |
| D114Y | Deficient | 0.9998479 | 0.978 |
| D122A | Deficient | 0.842692 | 0.941 |
| T125A | Deficient | 0.8518371 | 0.656 |
| C126W | Deficient | 0.9999958 | 0.996 |

| | | | |
|---|---|---|---|
| C126Y | Deficient | 1 | 0.997 |
| D127A | Deficient | 0.9998046 | 0.961 |
| R130I | Deficient | 0.9999819 | 0.993 |
| R130K | Deficient | 0.8419332 | 0.989 |
| L138R | Deficient | 0.8490442 | 0.509 |
| T142A | Proficient | 7.90496E-06 | 0.202 |
| Q146K | Proficient | 0 | 0.026 |
| Y148D | Deficient | 0.9867586 | 0.849 |
| D154A | Deficient | 0.7065448 | 0.388 |
| E159A | Proficient | 0.000376602 | 0.263 |
| E159G | Proficient | 0.000367283 | 0.221 |
| L162V | Proficient | 2.65583E-07 | 0.329 |
| F164C | Proficient | 0 | 0.047 |
| V166A | Proficient | 0 | 0.037 |
| K168N | Proficient | 7.04769E-07 | 0.062 |
| H172Y | Proficient | 0 | 0.036 |
| S173L | Proficient | 0 | 0.053 |
| Q185H | Proficient | 2.65583E-07 | 0.183 |
| V187A | Proficient | 0 | 0.099 |
| V187L | Proficient | 0 | 0.19 |
| S190A | Proficient | 0 | 0.071 |
| S190F | Proficient | 3.41371E-07 | 0.065 |
| L191V | Proficient | 0 | 0.009 |
| E198A | Proficient | 0 | 0.078 |
| A199V | Proficient | 0 | 0.047 |
| A203S | Proficient | 0 | 0.023 |
| E205A | Proficient | 0 | 0.035 |
| R207G | Deficient | 0.5102563 | 0.465 |
| R207Q | Proficient | 3.21269E-05 | 0.305 |
| R211Q | Proficient | 0 | 0.022 |
| E212K | Proficient | 0 | 0.022 |
| K213Q | Proficient | 0 | 0.068 |
| Q216K | Proficient | 0 | 0.04 |
| K217N | Proficient | 0 | 0.035 |
| K221E | Proficient | 0 | 0.06 |
| K222E | Proficient | 0 | 0.058 |
| R227W | Deficient | 0.649195 | 0.407 |
| S233N | Proficient | 0 | 0.046 |
| V234M | Proficient | 0 | 0.007 |
| T239M | Proficient | 0 | 0.007 |
| H242L | Deficient | 0.7065448 | 0.549 |
| E249Q | Proficient | 0 | 0.038 |
| R258C | Proficient | 0 | 0.055 |
| R258H | Proficient | 0 | 0.184 |
| T260N | Proficient | 0 | 0.011 |
| K272N | Proficient | 9.74785E-06 | 0.072 |

**Supplementary Table S6. Algorithm predictions for labeled XPA variants.**

| Variant | Training Label | Label Spreading Prediction | Label Spreading P(NER-deficient) | Logistic Regression Class Probability (NER-deficient) |
|---|---|---|---|---|
| K141E | Proficient | Proficient | 2.96719E-07 | 0.107 |
| K145E | Proficient | Proficient | 7.92005E-06 | 0.201 |
| K151E | Proficient | Proficient | 3.33862E-06 | 0.169 |
| C153S | Proficient | Proficient | 7.04769E-07 | 0.133 |
| K179E | Proficient | Proficient | 3.33862E-06 | 0.19 |
| K204E | Proficient | Proficient | 0 | 0.032 |
| R228Q | Proficient | Proficient | 0 | 0.054 |
| V234L | Proficient | Proficient | 0 | 0.051 |
| C105S | Deficient | Deficient | 0.9999999 | 0.996 |
| E106A | Deficient | Deficient | 0.9939312 | 0.835 |
| E106K | Deficient | Deficient | 0.9939312 | 0.809 |
| C108F | Deficient | Deficient | 1 | 0.997 |
| C108G | Deficient | Deficient | 1 | 0.997 |
| C108S | Deficient | Deficient | 0.9999999 | 0.996 |
| C126S | Deficient | Deficient | 0.9999999 | 0.996 |
| C129S | Deficient | Deficient | 0.9999999 | 0.996 |
| H244R | Deficient | Deficient | 0.7062171 | 0.635 |
| C261S | Deficient | Deficient | 0.9906542 | 0.92 |
| C264S | Deficient | Deficient | 0.9867586 | 0.887 |

**Supplementary Table S7. Average FM-HCR relative reporter expression normalized to wild-type compared with dbNSFP data for selected XPA variants.**

| | | Amino Acid Properties | Sequence Homology | | | Evolutionary Sequence Conservation | | | | | |
|---|---|---|---|---|---|---|---|---|---|---|---|
| Variant | FM-HCR Average Relative Reporter Expression (normalized to WT) *$p < 0.5$ | Grantham | SIFT | Polyphen2 | PROVEAN | LRT | MutationAssessor | GERP | phyloP | phastCons | ConSurf |
| D5Y | 1.03 | 160 | 0.043 | 0.101 | -1.12 | 0.800573 | 1.04 | 4.62 | 2.108 | 0.005 | 0.3643 |
| G6R | 1.03 | 125 | 0.54 | 0 | 0.23 | 0.122373 | 0.69 | -1.25 | 0.397 | 0 | 0.3427 |
| A18S | 1.16 | 99 | 1 | 0.001 | 0.78 | 0.038971 | 1.22 | 2.09 | 0.396 | 0 | 1.897 |
| R30W | 1.00 | 101 | 0 | 0.998 | -6.34 | 0 | 2.89 | 2.85 | 2.121 | 1 | -0.5574 |
| A60T | 1.02 | 58 | 0.745 | 0 | 1 | 0.048939 | 0.15 | 0.125 | -0.392 | 0.001 | 0.7473 |
| D70H | 1.01 | 81 | 0 | 0.999 | -6.86 | 0 | 3.055 | 5.45 | 4.593 | 1 | -1.035 |
| G72E | 1.04 | 98 | 0.045 | 0.762 | -6.98 | 0.001456 | 2.46 | 3.49 | 0.408 | 0.812 | -0.6186 |
| G73E | **0.49*** | 98 | 0.002 | 0.829 | -6.13 | 1E-06 | 2.9 | 4.55 | 4.593 | 1 | -0.7813 |
| P94L | 1.12 | 98 | 0.004 | 0.999 | -6.58 | 0 | 2.55 | 4.54 | 4.173 | 1 | 0.4415 |
| E106G | 1.08 | 98 | 0.366 | 0.021 | -3.66 | 0 | 0.75 | 5.25 | 5.491 | 1 | 1.165 |
| K110E | 1.02 | 56 | 0.183 | 0.265 | -1.61 | 0 | 1.1 | 5.25 | 7.195 | 1 | -0.06795 |
| E111A | 1.12 | 107 | 0.268 | 0.244 | -2.74 | 2.6E-05 | 1.245 | 5.25 | 4.636 | 1 | 2.09 |
| F112C | 0.96 | 205 | 0 | 0.997 | -7.1 | 0 | 2.215 | 5.25 | 8.38 | 1 | -0.6832 |
| M113I | **0.68*** | 10 | 0.202 | 0.71 | -1.69 | 0 | 1.65 | 5.25 | 7.096 | 1 | 0.1108 |
| D114Y | **0.72*** | 160 | 0.002 | 0.996 | -7.21 | 0 | 2.16 | 5.25 | 5.407 | 1 | -0.3485 |
| T125A | 1.14 | 58 | 0.071 | 0.111 | -0.76 | 8E-06 | 0.14 | 5.25 | 4.969 | 1 | -0.7623 |
| C126W | **0.07*** | 215 | 0 | 1 | -10.86 | 0 | 2.265 | 2.86 | 1.772 | 1 | -1.202 |
| C126Y | **0.09*** | 194 | 0 | 1 | -10.83 | 0 | 2.265 | 5.25 | 6.874 | 1 | -1.202 |
| R130I | 1.11 | 97 | 0.001 | 0.993 | -7.39 | 0 | 2.2 | 5.25 | 6.874 | 1 | -0.8641 |
| L138R | **0.72*** | 102 | 0 | 0.997 | -5.93 | 2.5E-05 | 3.325 | 5.34 | 8.891 | 1 | -0.9265 |
| Y148D | **0.50*** | 160 | 0 | 1 | -9.87 | 4E-06 | 3.28 | 5.34 | 8.891 | 1 | -0.6376 |
| D154A | 0.99 | 126 | 0 | 0.999 | -8 | 2E-06 | 3.045 | 5.34 | 7.628 | 1 | -1.174 |
| F164C | 1.06 | 205 | 0.148 | 0.181 | -5.65 | 7E-06 | 1.445 | 2.96 | 2.034 | 1 | 0.04784 |
| V234M | 0.95 | 21 | 0.213 | 0.027 | 0.11 | 0.067075 | 0.185 | 0.0447 | -0.24 | 0.009 | -0.2359 |
| H242L | 0.89 | 99 | 0 | 0.997 | -9.58 | 1E-06 | 2.97 | 5.32 | 7.072 | 1 | -1.209 |
| R258C | 0.99 | 180 | 0.137 | 0.424 | -1.33 | 0.05871 | 1.445 | 5.32 | 6.114 | 1 | 0.3136 |
| K272N | 0.95 | 94 | 0 | 0.994 | -4.51 | 0.002056 | 3.195 | 1.71 | 2.09 | 1 | -1.106 |

| | | Pathogenicity | | | | | Ensemble | | | |
|---|---|---|---|---|---|---|---|---|---|---|
| **Variant** | **FM-HCR Average Relative Reporter Expression (normalized to WT) *$p$ < 0.5** | **MutationTaster** | **FATHMM** | **VEST4** | **MPC** | **CADD** | **MetaSVM** | **MetaLR** | **REVEL** | **MVP** |
| **D5Y** | 1.03 | 1 | 0.13 | 0.3 | 0.325646623 | 1.973207 | -0.9851 | 0.1098 | 0.094 | 0.749718957 |
| **G6R** | 1.03 | 1 | 0.23 | 0.177 | 0.315686083 | 0.988008 | -1.0369 | 0.0943 | 0.052 | 0.512942373 |
| **A18S** | 1.16 | 0.999997 | 0.26 | 0.233 | 0.243273767 | 0.833877 | -1.0126 | 0.0776 | 0.044 | 0.538050723 |
| **R30W** | 1.00 | 1 | -0.14 | 0.717 | 0.703481252 | 4.74063 | -0.1566 | 0.4081 | 0.436 | 0.878378697 |
| **A60T** | 1.02 | 1 | 0.23 | 0.08 | 0.257212335 | 1.171349 | -1.092 | 0.064 | 0.062 | 0.485705204 |
| **D70H** | 1.01 | 1 | -0.55 | 0.702 | 1.027463522 | 3.890815 | 0.2192 | 0.5705 | 0.78 | 0.967765644 |
| **G72E** | 1.04 | 0.999996 | 0.1 | 0.253 | 0.589996054 | 2.765998 | -0.5521 | 0.2828 | 0.328 | 0.93410167 |
| **G73E** | **0.49*** | 0.999999 | -0.21 | 0.711 | 1.063991122 | 3.567701 | -0.0793 | 0.4271 | 0.522 | 0.932409099 |
| **P94L** | 1.12 | 1 | 0.17 | 0.854 | 0.411527654 | 4.005322 | -0.1844 | 0.3862 | 0.772 | 0.869043644 |
| **E106G** | 1.08 | 1 | -1.83 | 0.224 | 0.322851307 | 3.387624 | -0.402 | 0.438 | 0.339 | 0.942604788 |
| **K110E** | 1.02 | 1 | -2.17 | 0.489 | 0.386477495 | 3.303314 | 0.0089 | 0.5862 | 0.481 | 0.9335302 |
| **E111A** | 1.12 | 0.993409 | -1.7 | 0.211 | 0.318795421 | 3.177528 | -0.3251 | 0.4668 | 0.315 | 0.936076002 |
| **F112C** | 0.96 | 1 | -2.4 | 0.875 | 1.135461906 | 4.320504 | 0.8246 | 0.8225 | 0.928 | 0.97274872 |
| **M113I** | **0.68*** | 1 | -1.91 | 0.783 | 0.324661238 | 3.104572 | 0.1515 | 0.631 | 0.696 | 0.854444296 |
| **D114Y** | **0.72*** | 1 | -3.15 | 0.928 | 1.044414306 | 4.20561 | 0.9768 | 0.8987 | 0.875 | 0.984691032 |
| **T125A** | 1.14 | 0.734101 | -1.72 | 0.46 | 0.270821063 | 3.009224 | -0.3413 | 0.3444 | 0.293 | 0.893072399 |
| **C126W** | **0.07*** | 1 | -5.51 | 0.976 | 1.099507339 | 4.369788 | 1.092 | 0.9683 | 0.962 | 0.992237653 |
| **C126Y** | **0.09*** | 1 | -5.51 | 0.993 | 1.242852184 | 4.063909 | 1.0856 | 0.9766 | 0.909 | 0.983501949 |
| **R130I** | 1.11 | 1 | -4.91 | 0.834 | 1.256814846 | 3.996175 | 1.0764 | 0.9588 | 0.914 | 0.991642861 |
| **L138R** | **0.72*** | 1 | -0.66 | 0.986 | 1.146693974 | 4.685498 | 0.4037 | 0.6076 | 0.83 | 0.934173646 |
| **Y148D** | **0.50*** | 1 | -1.38 | 0.908 | 1.274480553 | 4.445982 | 0.6884 | 0.7396 | 0.91 | 0.966670494 |
| **D154A** | 0.99 | 1 | -0.23 | 0.851 | 0.965349088 | 4.247636 | 0.1982 | 0.5306 | 0.767 | 0.914097126 |
| **F164C** | 1.06 | 0.99999 | 0.14 | 0.335 | 0.418925122 | 3.040675 | -0.9411 | 0.1635 | 0.105 | 0.728141608 |
| **V234M** | 0.95 | 1 | 0.16 | 0.074 | 0.280752349 | 1.230607 | -1.0648 | 0.0819 | 0.019 | 0.248417906 |
| **H242L** | 0.89 | 1 | -0.55 | 0.79 | 1.051195728 | 4.003678 | 0.3237 | 0.5861 | 0.812 | 0.963922736 |
| **R258C** | 0.99 | 0.998891 | 0.14 | 0.374 | 0.772935469 | 4.147992 | -0.8437 | 0.1586 | 0.171 | 0.864606566 |
| **K272N** | 0.95 | 0.999598 | -0.37 | 0.663 | 1.018464281 | 4.170819 | -0.2552 | 0.4358 | 0.411 | 0.848901753 |

Where possible, cells are colored to reflect FM-HCR data or VEP tool predictions (pink for deleterious variants and green for tolerated variants).
* Signifies $p < 0.05$, unpaired t test.

**Supplementary Table S8. Average FM-HCR relative reporter expression normalized to wild-type compared with logistic regression model predictions for selected XPA VUS.**

| | Variant | Logistic Regression Class Probability P(NER-deficient) | FM-HCR Average Relative Reporter Expression (normalized to WT) **p* < 0.5 |
|---|---|---|---|
| Subset of Variants with Least Certain Class Probabilities | L138R | 0.509 | 0.72* |
| | H242L | 0.549 | 0.89 |
| | D70H | 0.418 | 1.01 |
| | E111A | 0.583 | 1.12 |
| | D154A | 0.388 | 0.99 |
| | T125A | 0.656 | 1.14 |
| | E106G | 0.663 | 1.08 |
| Variants for Evaluation | D5Y | 0.002 | 1.03 |
| | G6R | 0.022 | 1.03 |
| | A18S | 0.066 | 1.16 |
| | R30W | 0.098 | 1.00 |
| | A60T | 0.038 | 1.02 |
| | G72E | 0.06 | 1.04 |
| | G73E | 0.186 | 0.49* |
| | P94L | 0.208 | 1.12 |
| | K110E | 0.738 | 1.02 |
| | F112C | 0.962 | 0.96 |
| | M113I | 0.75 | 0.68* |
| | D114Y | 0.978 | 0.72* |
| | C126W | 0.996 | 0.07* |
| | C126Y | 0.997 | 0.09* |
| | R130I | 0.989 | 1.11 |
| | Y148D | 0.849 | 0.50* |
| | F164C | 0.047 | 1.06 |
| | V234M | 0.007 | 0.95 |
| | R258C | 0.055 | 0.99 |
| | K272N | 0.072 | 0.95 |

* Signifies $p < 0.05$, unpaired t test.
